# Linking YAP to Müller glia quiescence exit in the degenerative retina

**DOI:** 10.1101/431254

**Authors:** Annaïg Hamon, Divya Ail, Diana García-García, Juliette Bitard, Deniz Dalkara, Morgane Locker, Jérôme Roger, Muriel Perron

**Author notes:** These authors contributed equally to this work (Annaïg Hamon, Divya Ail, Diana García-García).

## Abstract

Contrasting with fish or amphibian, retinal regeneration from Müller glial cells is largely limited in mammals. In our quest towards the identification of molecular cues that may boost their stemness potential, we investigated the involvement of the Hippo pathway effector YAP, which we previously found to be upregulated in Müller cells following retinal injury. We report that conditional *Yap* deletion in Müller cells prevents the upregulation of cell cycle genes that normally accompanies reactive gliosis upon photoreceptor cell death. This occurs as a consequence of defective EGFR signaling. Consistent with a function of YAP in triggering Müller glia cell cycle re-entry, we further show that in *Xenopus*, a species endowed with efficient regenerative capacity, YAP is required for their injury-dependent proliferative response. Finally, and noteworthy, we reveal that YAP overactivation in mouse Müller cells is sufficient to induce their reprogramming into highly proliferative cells. Overall, we unravel a pivotal role for YAP in tuning Müller cell response to injury and highlight a novel YAP-EGFR axis by which Müller cells exit their quiescence state, a critical step towards regeneration.

## INTRODUCTION

Neurodegenerative retinal diseases, such as retinitis pigmentosa or age-related macular degeneration, ultimately lead to vision loss, as a consequence of photoreceptor cell death. Driving retinal self-repair from endogenous neural stem cells in patients represents an attractive therapeutic strategy. Among cellular sources of interest are Müller cells, the major glial cell type in the retina. These cells are essential for maintaining retinal homeostasis^1^ but also proved their potential for endogenous regeneration. In certain species, such as zebrafish or *Xenopus*, they behave as genuine stem cells, endowed with the ability to reprogram into a progenitor-like state upon retinal damage, proliferate and regenerate lost photoreceptors^2–4^. In mammals, however, their proliferative response to injury is extremely limited. Following acute retinal damage, mouse Müller glial cells rapidly re-enter G1-phase of the cell cycle, as inferred by increased *cyclin* gene expression, but they rarely divide ^5^. Suggesting that they nonetheless retain remnants of repair capacities, their proliferation and neurogenic potential can be stimulated, for instance by supplying exogenous growth factors such as Heparin-binding EGF-like growth factor (HB-EGF), or by overexpressing the proneural gene *Ascl1a*^3,6–8^. Our understanding of the genetic/signaling network sustaining Müller cell stemness potential is however far from being complete. Identifying novel molecular cues is thus of utmost importance to foresee putative candidates that could be targeted for regenerative medicine. We here investigated whether the Hippo pathway effector YAP might influence Müller cell reactivation and how it would intersect with other critical signaling pathways.

The Hippo pathway is a kinase cascade that converges towards two terminal effectors, YAP (Yes-associated protein) and TAZ (transcriptional coactivator with PDZ-binding motif). Both are transcriptional coactivators of TEAD family transcription factors. Originally identified in *Drosophila* as a regulator of organ growth, the Hippo pathway progressively emerged as a key signaling in a wide range of biological processes^9^, including stem cell biology^10,11^. Of note, it proved to be dispensable under physiological conditions in some adult stem cells, such as mammary gland, pancreatic, intestinal and, importantly, neural stem cells^12–15^. It can nonetheless become essential under pathological conditions, as described for example in the context of intestinal regeneration following injury^16,17^. YAP status in adult neural tissue repair has hitherto never been investigated. We recently discovered that YAP and TEAD1 are specifically expressed in murine Müller cells and that their expression and activity are enhanced upon retinal damage^18^. We thus sought to determine whether YAP could be required for injury-induced Müller glia reactivation. We found in mouse that YAP triggers cell cycle gene upregulation in Müller glial cells following photoreceptor cell death, a function that likely relies on its interplay with EGFR signaling. In line with the idea of a conserved role in Müller cell cycle re-entry, blocking YAP function in *Xenopus* results in a dramatically reduced proliferative response following acute retinal damage or photoreceptor cell ablation. Finally, we report that the limited proliferative response of murine Müller glia can be circumvented and significantly enhanced by YAP overexpression. As a whole, this study highlights the critical role of YAP in driving Müller cells to exit quiescence and thus reveals a novel potential target for regenerative medicine.

## RESULTS

### *Yap* conditional knockout in mouse Müller cells does not compromise their maintenance under physiological conditions

To investigate the role of YAP in murine Müller glia, we generated a *Yap^flox/flox^; CreER^T2^* mouse line allowing for Cre-mediated conditional gene ablation specifically in Müller cells^19,20^. It is thereafter named *Yap* CKO while “control” refers to *Yap*^flox/flox^ mice. *Yap* deletion was induced in fully differentiated Müller cells, through 4-hydroxytamoxifen (4-OHT) intraperitoneal injection at P10 (Fig. 1A). Phenotypic analyses were then conducted on 2-month-old mice. We first confirmed Müller cell-specific *Cre* expression and *Yap* deletion efficiency in *Yap* CKO carrying the Rosa26-CAG-lox-stop-lox-TdTomato reporter (*Ai9*) transgene (Fig. 1B, C). We next wondered whether expression of TAZ (the second effector of the Hippo pathway) could be increased in our model and thus potentially compensate for *Yap* deletion, as previously reported in mammalian cell lines^21^. In physiological conditions, TAZ protein level is actually similar in *Yap* CKO and control mice (Supplementary Figure S1A), suggesting an absence of compensatory mechanisms. Finally, we assessed global retinal organization and function in *Yap* CKO mice. Immunostaining for various retinal neuron markers and electroretinogram (ERG) recordings under scotopic and photopic conditions revealed no structural nor functional difference compared to control retinas (Supplementary Figure S1B-D). Müller cells were also normally distributed and did not display any sign of stress reactivity, as assessed by the expression of intermediate filament proteins, glial fibrillary acidic protein (GFAP) and Vimentin (Fig. 1D, E). Taken together, these results demonstrate that lack of YAP expression in Müller cells from P10 does not impact the overall retinal structure, neuron and glia maintenance, nor the visual function in 2-month-old mice.

**Figure 1:**
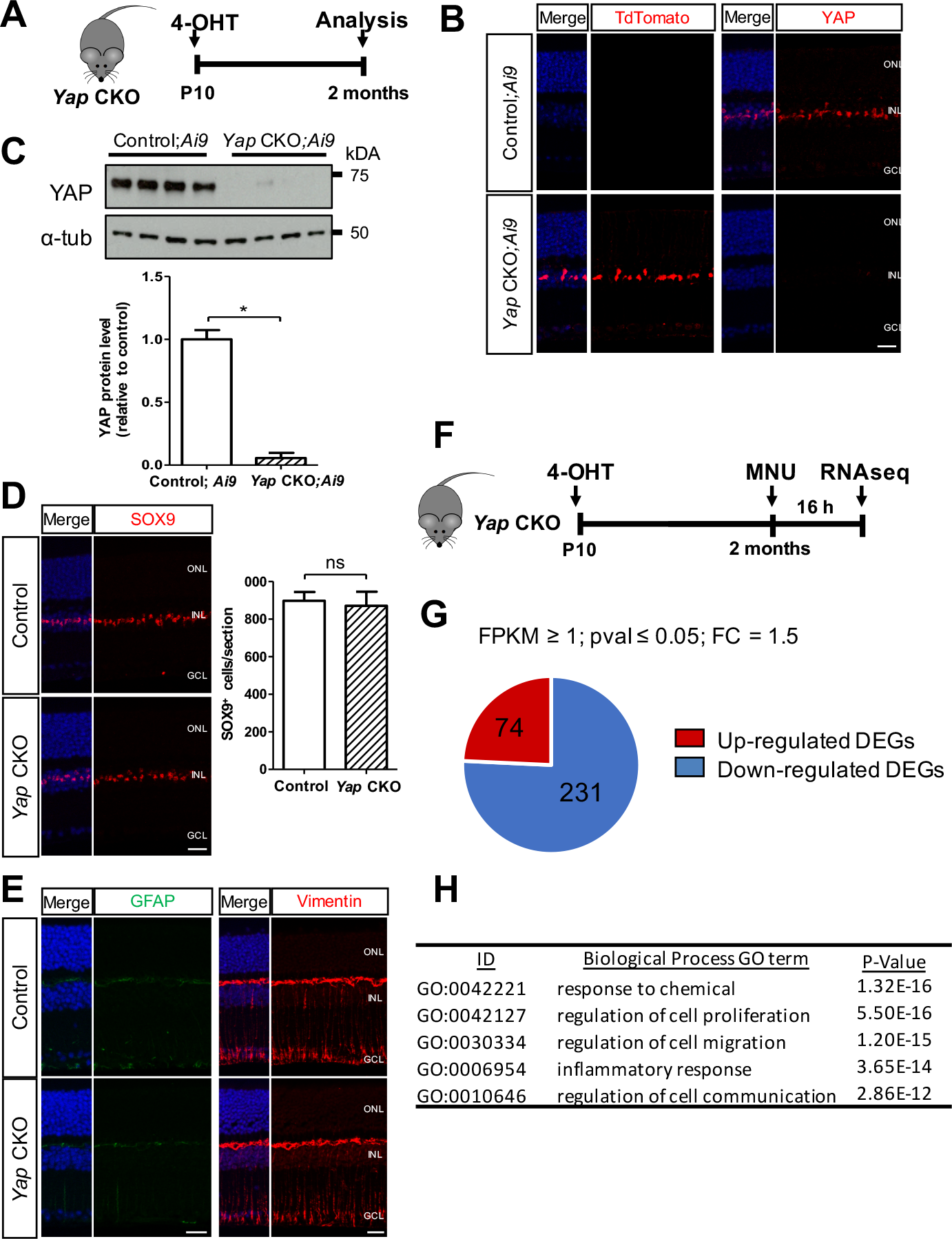
*Yap* CKO Müller cells exhibit altered transcriptional response to injury.. **(A)** Timeline diagram of the experimental procedure used in B-E. *Yap*^*flox/flox*^ mice with or without the *Ai9* reporter allele (Control or Control;/*Ai9*) and *Yap*^*flox/flox*^;*Rax-CreER*^*T2*^ mice with or without the *Ai9* reporter allele (*Yap* CKO or *Yap* CKO;/*Ai9*) received a single dose of 4-OHT at P10 and were analyzed at P60. **(B)** Retinal sections immunostained for TdTomato or YAP. **(C)** Western-blot analysis of YAP expression on retinal extracts, *α*-tubulin labelling was used to normalize the signal. Quantification: n=4 mice for each condition. **(D)** Retinal sections immunostained for SOX9. Quantification: n=3 mice for each condition. **(E)** Retinal sections immunostained for GFAP or Vimentin. **(F)** Timeline diagram of the experimental procedure used in G and H. *Yap*^*flox/flox*^ (Control) and *Yap*^*flox/flox*^;*Rax-CreER*^*T2*^ (*Yap* CKO) mice received a single dose of 4-OHT at P10 and a single dose of MNU at 2 months. Retinas were then subjected to RNA-sequencing 16 hours later. **(G)** Pie chart representing the number of DEGs found to be up- or downregulated in MNU-injected *Yap* CKO retinas compared to MNU-injected control ones. **(H)** Results of Gene Ontology (GO) enrichment analysis exemplifying six over-represented GO biological processes related to retinal response to injury (the complete top twenty can be found in Supplementary Figure S3). ONL: outer nuclear layer; INL: inner nuclear layer; GCL: ganglion cell layer. In (B, D, E), nuclei were counterstained with DAPI (blue). Statistics: Mann-Whitney test, *p≤ 0.05, ns: non-significant. Scale bars: 20 μm.

### *Yap* deletion impairs mouse Müller cell reactivation upon photoreceptor degeneration

We next investigated YAP function in a degenerative context. Retinal degeneration was triggered in *Yap* CKO mice through *in vivo* methylnitrosourea (MNU) injections, a well-established paradigm for inducible photoreceptor cell death^22^. In order to evaluate the impact of *Yap* deletion on Müller glia early response to injury, all the analyses were performed 16 hours after MNU injection, at the onset of photoreceptor cell death (Fig. 1F and Supplementary Figure S2A). As previously described^18^, YAP protein levels were upregulated in WT retinas upon MNU injection and, as expected, we found them effectively decreased in our MNU-injected *Yap* CKO model (Supplementary Figure S2B). In contrast with the physiological situation, this was accompanied by a compensatory increase of TAZ levels (Supplementary Figure S2C). Yet, this enhanced expression is likely insufficient to entirely compensate for the loss of YAP activity, as inferred by the downregulation of *Cyr61*, a well-known YAP/TAZ target gene^23^ (Supplementary Figure S2D). We next assessed GFAP expression as a marker of reactive gliosis upon MNU injection. Interestingly, we found it increased at both the mRNA and protein levels in *Yap* CKO mice compared to control ones, reflecting a potential higher degree of retinal stress (Supplementary Figure S2E, F). This led us to further investigate the molecular impact of *Yap* deletion in reactive Müller cells, through a large-scale transcriptomic analysis comparing non-injected WT mice, MNU-injected control mice and MNU-injected *Yap* CKO mice. This allowed identifying 305 differentially expressed genes (DEGs), 75% of them being downregulated in MNU-injected *Yap* CKO retinas compared to MNU-injected control ones (Fig. 1G). The top-enriched biological processes they belong to include “response to chemical”, “regulation of cell proliferation” and “inflammatory response”. This strongly suggests that lack of YAP expression profoundly alters Müller cell transcriptional response to retinal injury (Fig. 1H and Supplementary Figure S3).

### *Yap* knockout prevents cell cycle gene up-regulation in mouse reactive Müller cells

As we were seeking for a potential function of *Yap* in Müller cell reactivation, we focused our interest on identified genes related to the GO group “regulation of cell proliferation”. Z-score-based hierarchical clustering for the 70 corresponding DEGs revealed 3 distinct clusters (Supplementary Figure S4). One particularly caught our attention since the 52 DEGs it contains appear less responsive to injury in the absence of YAP. These are indeed (*i*) expressed at very low levels in wild type mice, (*ii*) strongly upregulated in MNU-injected control mice, (*iii*) while being only moderately enriched in MNU-injected *Yap* CKO mice. This is the case for instance of four cell cycle regulator coding genes, *Ccnd1, Ccnd2, Ccnd3* and *Cdk6* (Fig. 2A). Further validation was conducted by qPCR and western-blot for Cyclin D1 and Cyclin D3, which are specifically expressed in Müller cells. This confirmed their downregulation upon MNU injection in *Yap* CKO mice compared to controls (Fig. 2B-D). Noticeably, we found that the pluripotent factor LIF and the reprogramming factor Klf4 follow the same profile, suggesting that the reprogramming process that initiates along with Müller cell reactive gliosis is also impaired by *Yap* loss of function (Supplementary Figure S5).

**Figure 2:**
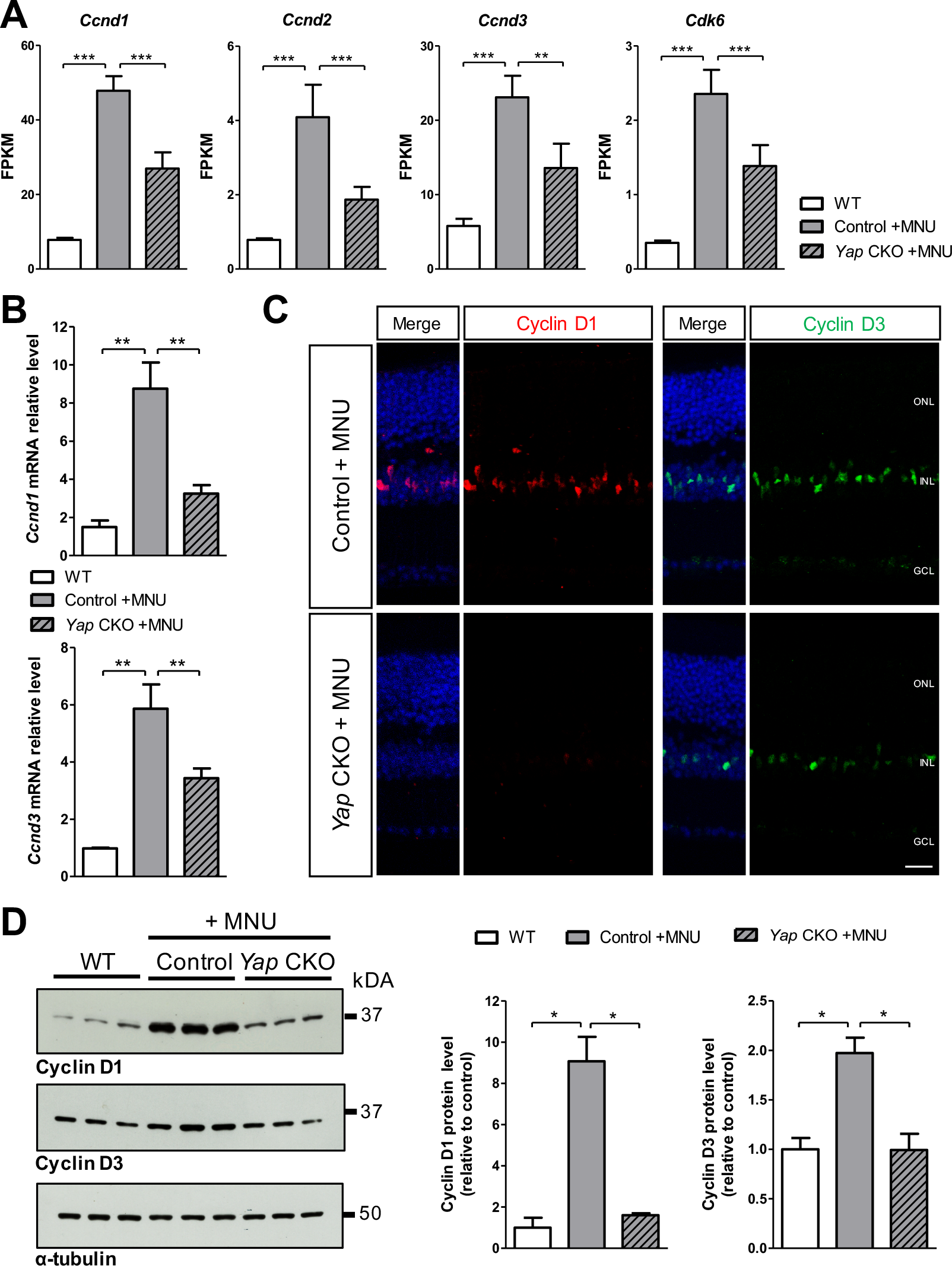
Cell cycle gene upregulation in response to MNU injection is compromised in *Yap* CKO retinas.. **(A)** Relative RNA expression (in FPKM; data retrieved from the RNA-seq experiment) of *Ccnd1*, *Ccnd2*, *Ccnd3* and *Cdk6*, in retinas from non-injected WT mice or Control and *Yap* CKO mice injected with 4-OHT and MNU as shown in Figure 1F. **(B)** RT-qPCR analysis of *Ccnd1* and *Ccnd3* expression in the same experimental conditions (at least five biological replicates per condition were used). **(C)** Retinal sections from Control and *Yap* CKO mice, immunostained for Cyclin D1 or D3. Nuclei are counterstained with DAPI (blue). **(D)** Western-blot analysis of Cyclin D1 and D3 expression on retinal extracts from WT, Control and *Yap* CKO mice, *α*-tubulin labelling was used to normalize the signal. Quantification: n=3 mice for each condition. ONL: outer nuclear layer; INL: inner nuclear layer; GCL: ganglion cell layer. Statistics: Mann-Whitney test (except in A where P-values were obtained using EdgeR), *p≤ 0.05, **p≤ 0.01; ***p≤ 0.001. Scale bar: 20 μm.

To strengthen our results in a model closer to human retinal disease, we bred the *Yap* CKO line into the *rd10* background (*Pde6b*^*rd10*^ line, a model of retinitis pigmentosa^24,25^). We next assessed Cyclin D1 and Cyclin D3 expression at P20 (Fig. 3A, B), which corresponds to the period of intense rod cell death in *rd10* mice^24^. As observed with the MNU model, labelling for both cyclins was increased in Müller cells upon photoreceptor degeneration, and this upregulation was impaired in *Yap* CKO retinas. Taken together, these results demonstrate YAP requirement for the injury-induced transcriptional activation of cell cycle genes, which likely reflects its role in pushing Müller cells out of their quiescent state. Of note, a YAP-dependent control of cyclin D1 and D3 expression was also observed in physiological conditions (Fig. 3C, D), suggesting that YAP regulates the basal level of cell cycle genes in quiescent Müller cells as well.

**Figure 3:**
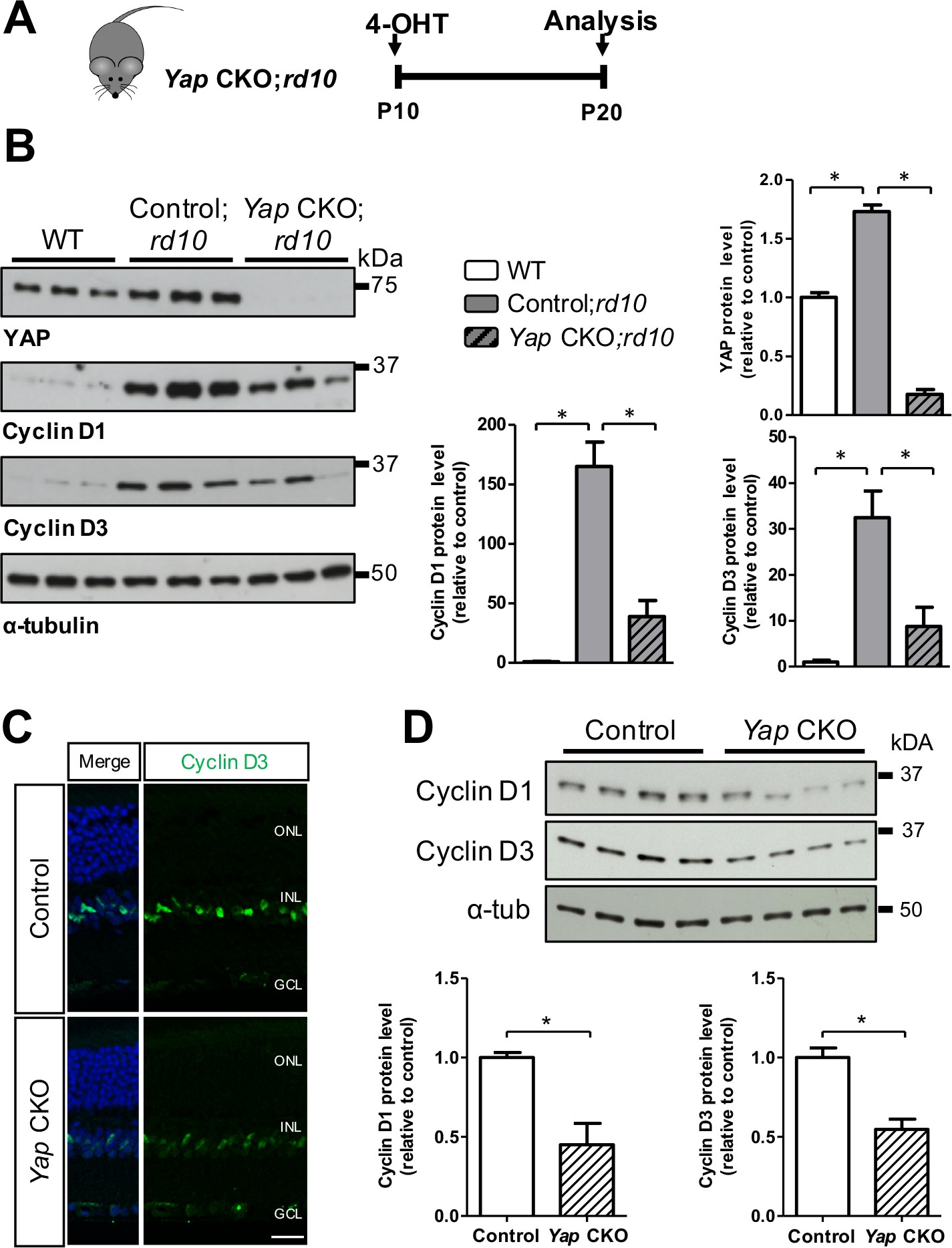
YAP loss of function results in decreased Cyclin D1/D3 expression in *rd10* degenerating retinas or in a physiological context.. (**A**) Timeline diagram of the experimental procedure used in B. *Yap*^*flox/flox*^;*rd10* (Control;*rd10*) and *Yap*^*flox/flox*^;*Rax-CreER*^*T2*^;*rd10* (*Yap CKO;rd10*) mice received a single dose of 4-OHT at P10 and were analyzed at P20. (**B**) Western-blot analysis of YAP, Cyclin D1 and Cyclin D3 expression, *α*-tubulin labelling was used to normalize the signal. Quantification: n=3 mice for each condition. **(C)** Retinal sections from 2-month-old Control and *Yap* CKO mice in physiological conditions, immunostained for Cyclin D3. Nuclei are counterstained with DAPI (blue). **(D)** Western-blot analysis of Cyclin D1 and Cyclin D3 expression on retinal extracts from 2-month-old Control and *Yap* CKO mice in physiological conditions, α-tubulin labelling was used to normalize the signal. Quantification: n=4 mice for each condition. ONL: outer nuclear layer; INL: inner nuclear layer; GCL: ganglion cell layer. Statistics: Mann-Whitney test, *p≤ 0.05. Scale bar: 20 μn.

### *Yap* deletion affects EGFR signaling in mouse reactive Müller cells

Beside cell cycle genes, our RNAseq dataset also revealed a deregulation of several members of the EGFR (Epidermal Growth Factor Receptor) pathway in the *Yap* CKO degenerative background. Importantly, EGFR signaling is well known for its mitogenic effects on Müller cells during retinal degeneration. Two EGFR ligands, namely EGF or HB-EGF, have in particular been shown to stimulate Müller glia proliferation in zebrafish, chick or rodents^26–30^. As observed with cell cycle genes, both *Egfr* and two ligand-coding genes (*Hbegf* and *Neuregulin 1*) failed to be properly upregulated upon MNU-injection in *Yap* CKO retinas compared to control ones (Fig. 4A). Expression of another EGFR-coding gene, *Erbb4* (named also *Her4*), did not appear sensitive to MNU injection in control retinas but was still found significantly decreased in MNU-injected *Yap* CKO mice. In order to decipher whether these deregulations might be associated with defective EGFR signaling activity, we next assessed the status of the MAPK and the PI3K/AKT pathways, which are known to be required for Müller cell proliferative response to growth factor treatment upon injury^31–33^. Western-blot analysis of ERK1/2 and AKT phosphorylation revealed increased P-ERK/ERK and P-AKT/AKT ratios following MNU injection in control retinas. In the *Yap* CKO context, this increase was significantly attenuated, reflecting lower signaling activation (Fig. 4B). Importantly, P-ERK immunostaining confirmed that this took indeed place in Müller cells: P-ERK labelling (which is barely detectable in non-injured retinas; data not shown) was localized in Müller cell nuclei and processes upon MNU injection and exhibited differential enhancement in control mice (strong signal) compared to *Yap* CKO ones (weaker signal; Fig. 4C). Together, these results suggest that YAP is required for proper EGFR signaling through its transcriptional control on both EGFR ligands and receptors. Of note, pathway analysis from our RNAseq dataset revealed that other genes belonging to the PI3K-AKT and ErbB signaling pathways were also deregulated in *Yap* CKO retinas, with 15 and 6 associated DEGs, respectively (Supplementary Figure S6). This further supports the existence of a functional link between YAP and the EGFR pathway in the degenerating retina.

**Figure 4:**
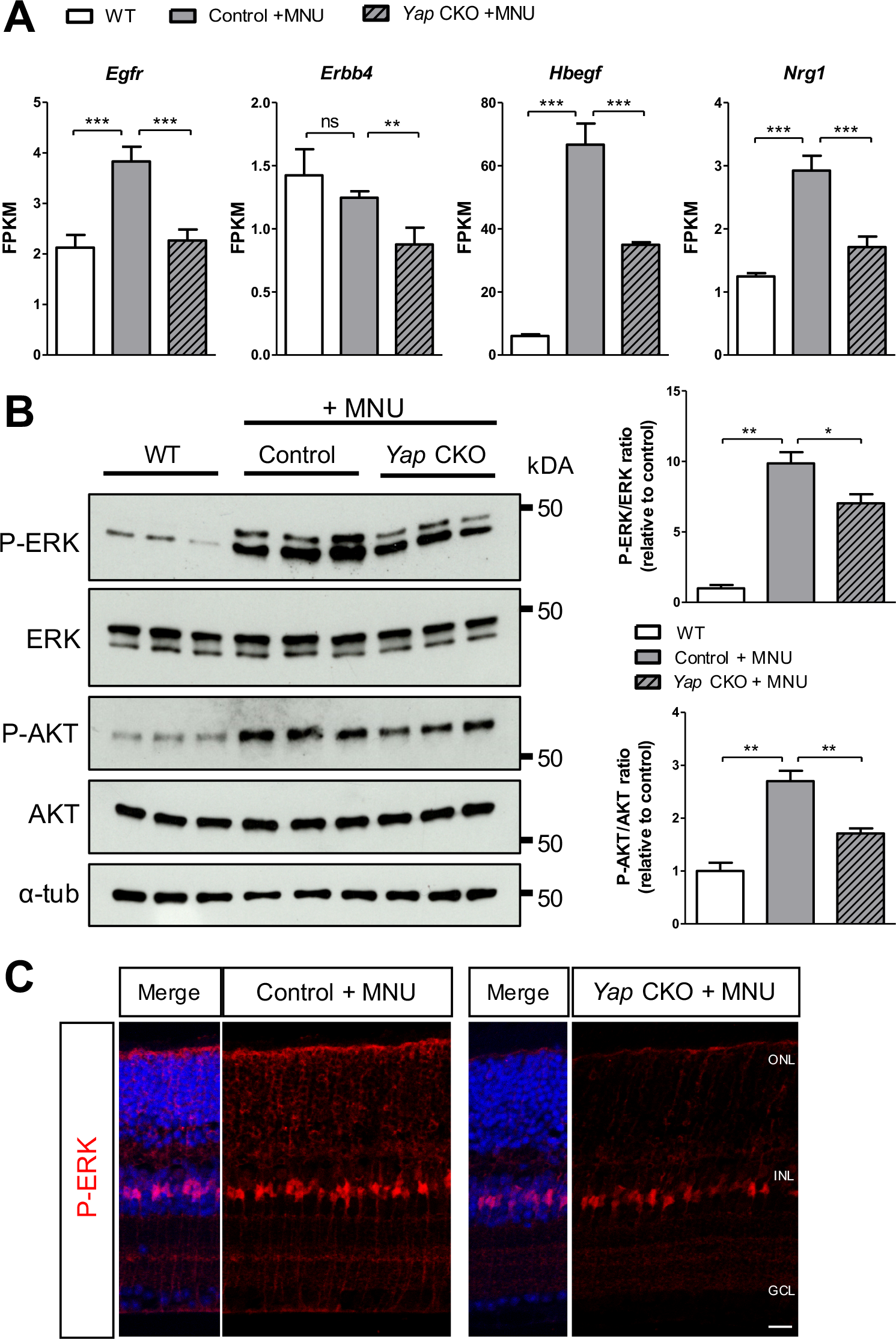
*Yap* CKO Müller cells display altered EGFR signaling in response to injury.. **(A)** Relative RNA expression (in FPKM; data retrieved from the RNA-seq experiment) of *Egfr*, *Erbb4*, *Hbegf* and *Neuregulinl* (*Nrg1*), in retinas from non-injected WT mice or Control and *Yap* CKO mice injected with 4-OHT and MNU as shown in Fig. 1F. **(B)** Western-blot analysis of P-ERK/ERK and P-AKT/AKT ratios on the same experimental conditions, *α*-tubulin labelling was used to normalize the signal. Quantification: n=6 mice for each condition. **(C)** Retinal sections from Control and *Yap* CKO mice, immunostained for P-ERK. Nuclei are counterstained with DAPI (blue). ONL: outer nuclear layer; INL: inner nuclear layer; GCL: ganglion cell layer. Statistics: Mann-Whitney test (except in A where P-values were obtained using EdgeR), *p≤ 0.05, **p≤ 0.01; ***p≤ 0.001, ns: non-significant. Scale bar: 20 μm.

### HB-EGF treatment rescues cell cycle gene expression in *Yap* CKO mouse reactive Müller cells

We next sought to investigate whether this YAP-EGFR interaction might converge on cell cycle gene regulation, by attempting to rescue the *Yap* CKO phenotype through EGFR signaling activation. To this aim, we decided to treat *Yap* CKO retinas with HB-EGF, one of the EGFR ligands that we found decreased in the absence of YAP (Fig. 4A). We performed the experiment *ex vivo* on adult retinal explants, a spontaneous model of retinal degeneration that facilitates the use of exogenous factors^34^. As expected from our previous observations in MNU and *rd10* mice^18^, we found an increase in YAP level accompanying the degenerative process in explants (Fig. 5A, B). Moreover, as described above with both paradigms (Figs. 2 and 3), this correlated with Cyclin D1 upregulation (Fig. 5D; compare lanes 1 and 2), and this response was impaired in *Yap* CKO explants (Fig. 5D, compare lanes 2 and 3). Remarkably, following HB-EGF addition, Cyclin D1 levels were indistinguishable between *Yap* CKO and control explants (Fig. 5C, D). Such a rescue of *Yap* deletion by exogenous supply of HB-EGF strongly suggests that YAP is acting upstream the EGFR pathway in Müller glia cell cycle gene regulation.

**Figure 5:**
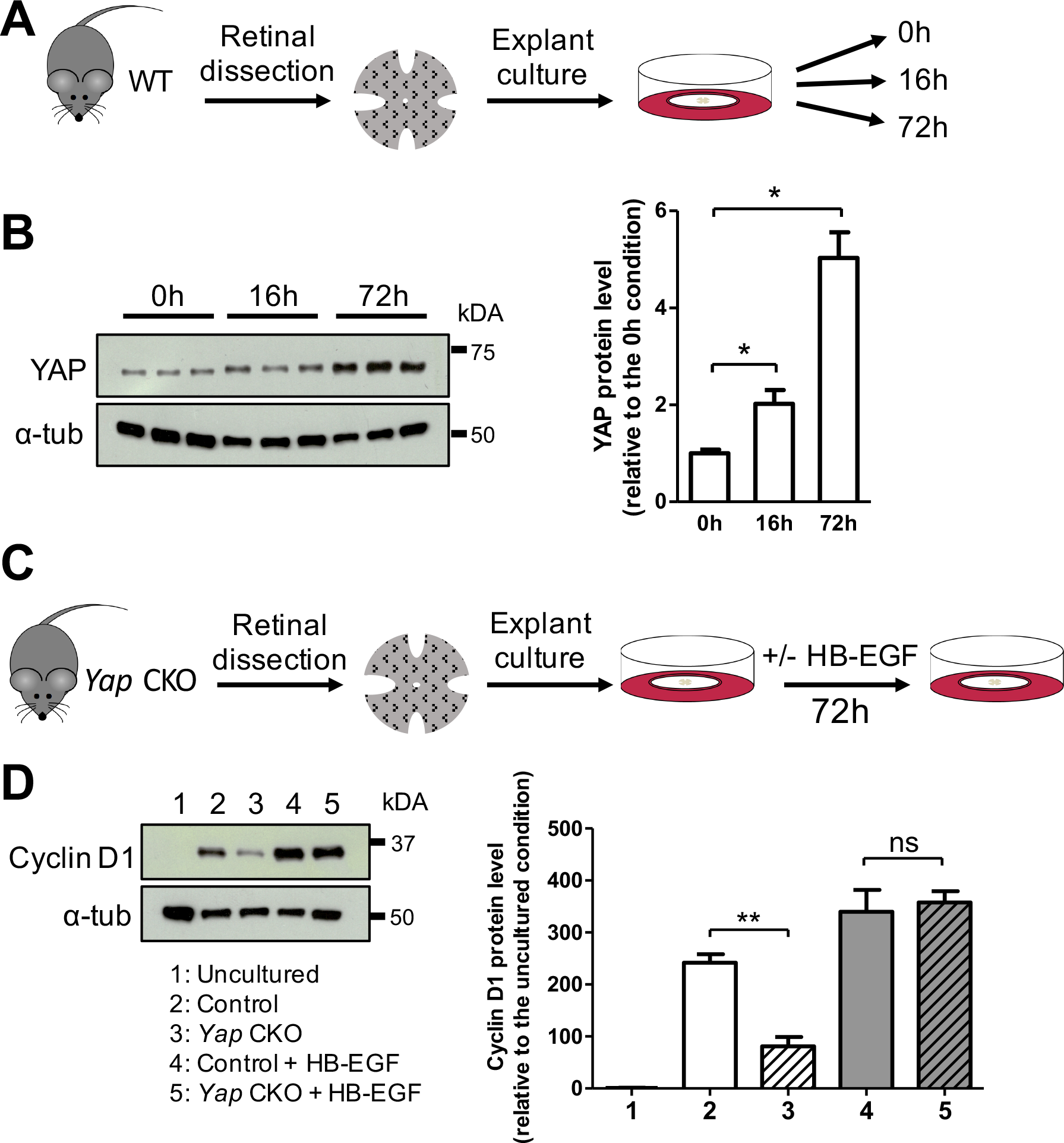
YAP CKO phenotype can be rescued by HB-EGF treatment.. **(A)** Timeline diagram of the experimental procedure used in B. Retinas from WT mice were flattened and cultured for 0, 16 or 72 hours. **(B)** Western-blot analysis of YAP expression on retinal explant extracts, α-tubulin labelling was used to normalize the signal. Quantification: n=3 mice for each condition. **(C)** Timeline diagram of the experimental procedure used in D. Retinas from Control and *Yap* CKO mice were cultured for 72 hours with or without HB-EGF. **(D)** Western-blot analysis of Cyclin D1 expression on retinal explant extracts, α-tubulin labelling was used to normalize the signal. Quantification: n=3 explants for the uncultured condition, n=6 explants for all other conditions. Statistics: Mann-Whitney test, *p≤ 0.05, **p≤ 0.01; ns: non-significant.

### Inhibition of YAP prevents Müller glia proliferation upon acute retinal damage or selective photoreceptor cell ablation in *Xenopus laevis*

All the above results converge to the idea that YAP triggers cell cycle re-entry of quiescent Müller glia upon injury. Since this process is not complete in murine Müller cells (they reactivate G1-phase genes but rarely divide), we turned to the frog to strengthen our hypothesis. *Xenopus* is an animal model endowed with regenerative capacity, in which Müller cells efficiently respond to injury by intense proliferation^4^. We first confirmed that, within the *Xenopus* central neural retina, YAP expression is restricted to Müller cells (Fig. 6A and ^35^), as observed in mouse. We next sought to assess the impact of YAP inhibition by taking advantage of a *Xenopus laevis* transgenic line, hereafter named *Tg(dnYAP)*, in which a heat shock promoter (Hsp70) drives the expression of a dominant-negative YAP variant^36,37^ (Fig. 6B and Supplementary Figure S7A, B). Confirming the transgene efficiency, two YAP target genes, *Ctgf* (connective tissue growth factor) and *Cyr61*, were downregulated in *Xenopus Tg(dnYAP)* retinas upon heat-shock induction (Supplementary Figure S7C). We next assayed Müller cells proliferative response at tadpole stage in a model of stab injury (Fig. 6C). Importantly, we previously demonstrated that a majority of proliferating cells found at the injury site are indeed Müller cells^4^. Comparison of BrdU incorporation in heat-shocked and non-heat-shocked wild-type retinas, confirmed that heat-shock does not affect proliferation by itself (Fig. 6D). In contrast, the number of BrdU-labelled cells at the injury site was reduced by about 60% in heat-shocked *Tg(dnYAP)* tadpole retinas compared to controls (non-heat-shocked transgenic animals or heat-shocked non-transgenic siblings; Fig. 6D). Of note, loss of YAP activity did not affect Müller cell number (Supplementary Figure S7D), ruling out the possibility that defective BrdU incorporation might be due to an impaired cell survival. Finally, YAP requirement for Müller glia proliferative response to stab injury could be confirmed at post-metamorphic stage in froglets, with *Tg(dnYAP)* retinas exhibiting a 80% reduction of BrdU-positive cells compared to controls (Fig. 6E, F).

**Figure 6:**
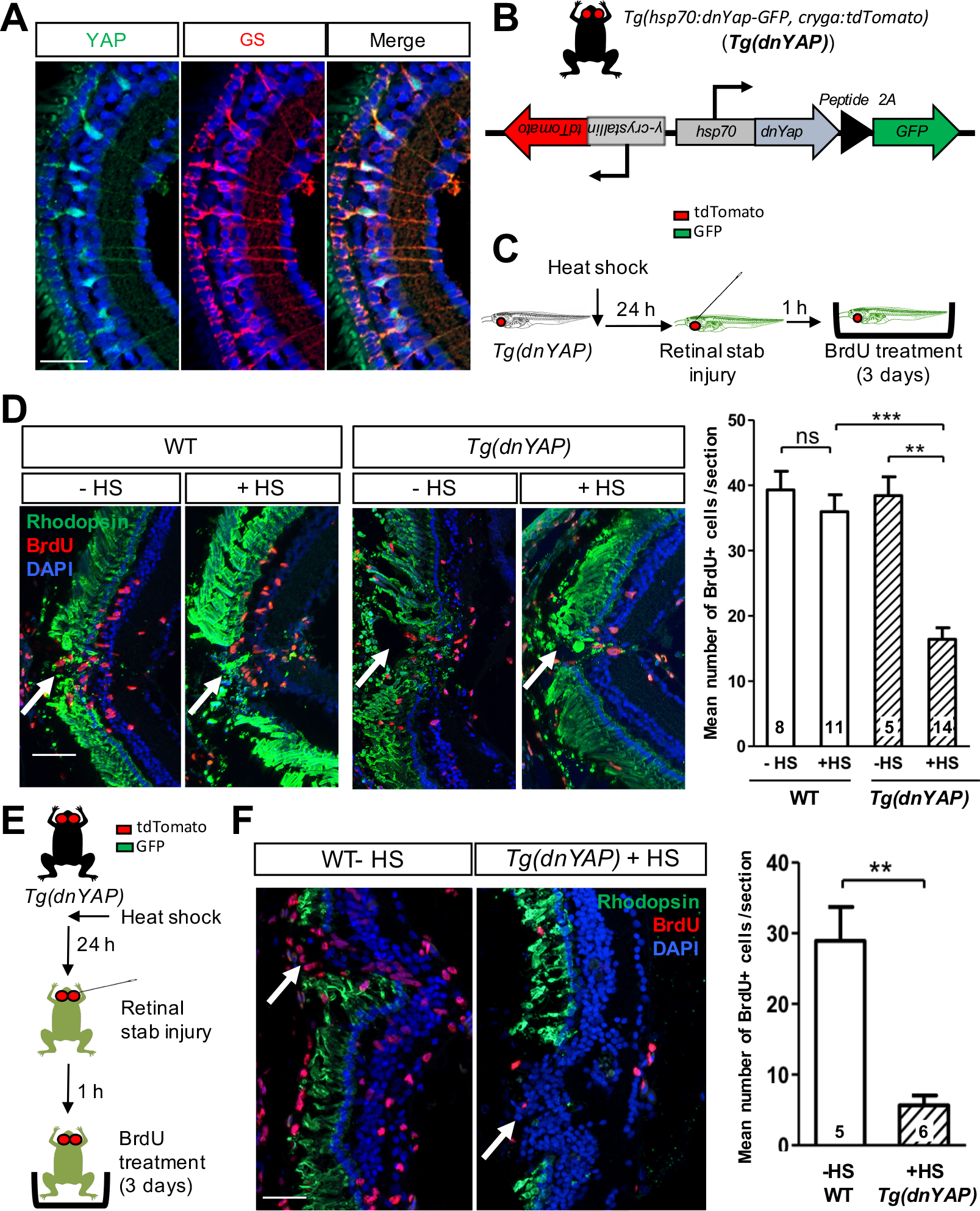
Inhibiting YAP activity in *Xenopus* reduces the proliferative retinal response to a stab injury.. (**A**) Retinal section from a stage 45 WT *Xenopus laevis* tadpole, immunostained for YAP and Glutamine synthetase (GS), a marker of Müller cells. Nuclei are counterstained with Hoechst (blue). (**B**) Schematic representation of the heat-shock inducible dominant-negative *Yap* transgene (*dnYap*). *Tg(dnYAP)* transgenic animals can be selected based on tdTomato expression in the lens (driven by the (-crystallin promoter). Heat-shocked efficiency can be assessed through GFP expression (see Supplementary Figure S7B). (**C, E**) Timeline diagrams of the experimental procedures used in D and F, respectively. WT or *Tg(dnYAP)* pre-metamorphic tadpoles (stage 54-58, D) or froglets (stage 61-66, F) were heat-shocked, stabbed injured in the retina 24 hour later, and transferred one hour post-lesion in a BrdU solution for 3 more days. (**D, F**) Retinal sections from animals that were heat-shocked (+HS) or not (-HS), immunostained for Rhodopsin (green) and BrdU. Nuclei are counterstained with DAPI (blue). Arrows point to the injury site. Quantification: the number of animals tested for each condition is indicated on the corresponding bar. Statistics: Mann-Whitney test, **p≤ 0.01, ***p≤ 0.001; ns: non-significant. Scale bars: 25 μm (A), 50 μm (D, F).

We next sought to reinforce these data in a model closer to the mouse MNU or *rd10* paradigms. In this purpose, we turned to a *Xenopus laevis* transgenic line that we previously established, allowing for conditional selective rod cell ablation^4^ (*Tg(Rho:GFP-NTR)*, hereafter named *Tg(NTR)*; Fig. 7A). This transgenic line expresses the nitroreductase (*NTR*) gene under the control of the *Rhodopsin* promoter and photoreceptor degeneration can be induced by adding the enzyme ligand metronidazole (MTZ) to the tadpole rearing medium. Here again, we previously showed that about 80% of cells that proliferate upon rod cell ablation are indeed Müller cells^4^. The *Tg(dnYAP)* and *Tg(NTR)* lines were crossed to generate double transgenic animals, in which inhibition of YAP can be triggered by heat-shock, and photoreceptor degeneration by MTZ treatment (Fig. 7B). As observed above with the stab injury, Müller cell proliferative response to rod cell death was dramatically reduced in double *Tg(dnYAP;NTR)* froglet retinas compared to *Tg(NTR)* control ones (Fig. 7C). Altogether, this supports the idea that YAP is required for *Xenopus* Müller glia cell cycle re-entry and proliferation, in different lesional contexts.

**Figure 7:**
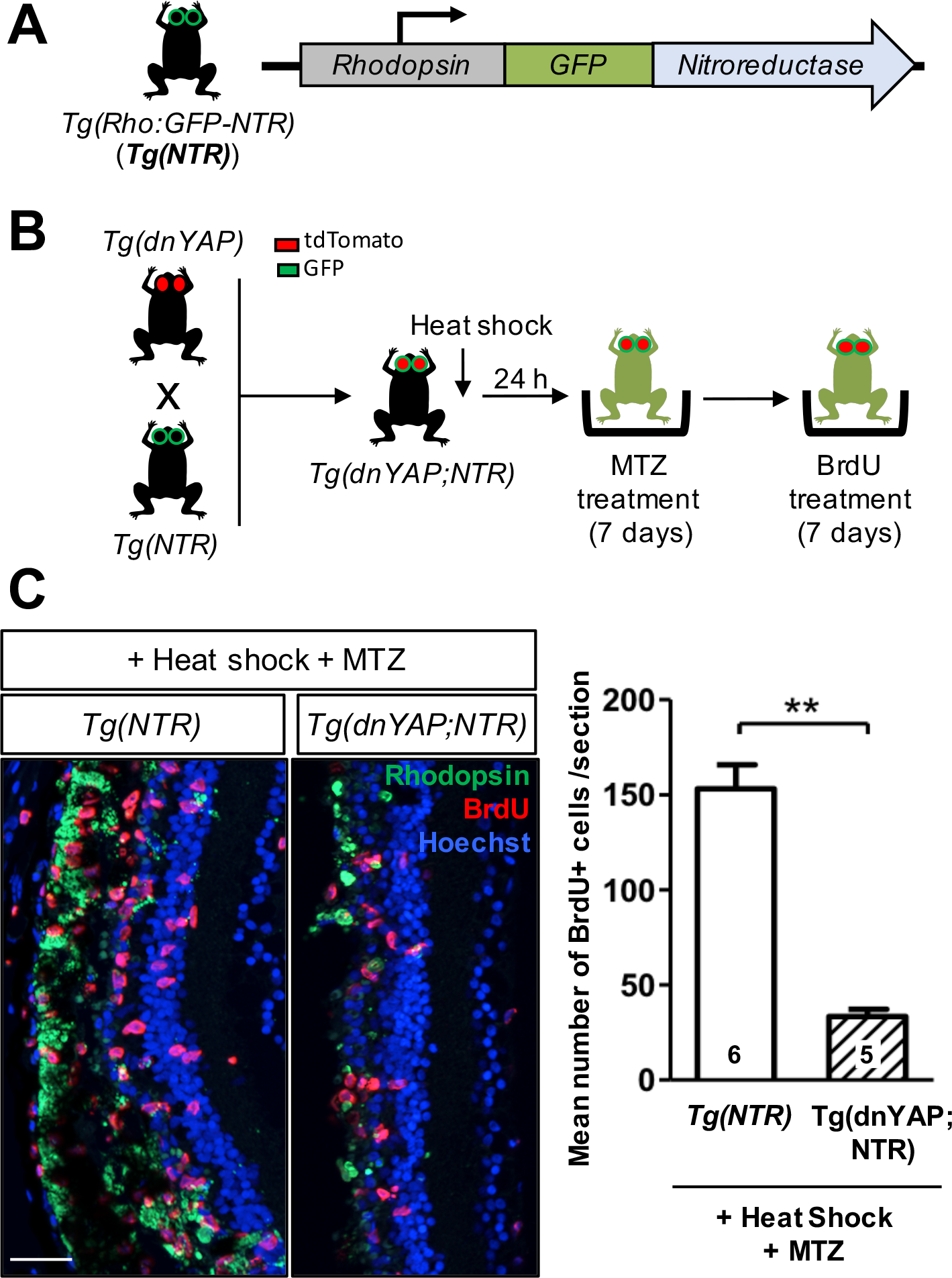
Inhibiting YAP activity in *Xenopus* reduces the proliferative retinal response to photoreceptor degeneration.. (**A**) Schematic representation of the transgene carried by the *Tg(NTR)* line. Transgenic animals can be selected based on GFP expression in rod cells (driven by the *Rhodopsin* promoter). (**B**) Timeline diagram of the experimental procedures used in C. *Tg(dnYAP)* were crossed with *Tg(NTR)* to generate double transgenic *Tg(dnYAP;NTR)* animals. *Tg(dnYAP;NTR)* froglets (stage 61-66) were heat-shocked, transferred 24 hours later for seven days in a MTZ solution, and finally exposed for seven more days to BrdU. (**C**) Retinal sections from control (*Tg(NTR)*) and *Tg(dnYAP;NTR)* animals, immunostained for Rhodopsin and BrdU. Nuclei are counterstained with DAPI (blue). Note the scattered green staining indicative of photoreceptor degeneration. Quantification: the number of animals tested for each condition is indicated on the corresponding bar. Statistics: Mann-Whitney test, **p≤ 0.01. Scale bars: 50 μm.

### Forced YAP expression in mouse Müller glia cells stimulates their proliferation in retinal explants

Based on the above data on *Xenopus*, we next wondered whether mouse Müller cell inability to proliferate upon injury (despite cell cycle gene reactivation), might be linked to insufficient levels of YAP activity. To investigate this hypothesis, we sought to overexpress a mutated YAP protein, YAP^5SA^, which is insensitive to Hippo pathway-mediated cytoplasmic retention^38^. To deliver this constitutively active form of YAP, we infected retinal explants with an adeno-associated virus (AAV) variant, ShH10, which selectively targets Müller cells^39^ (Fig. 8A and Supplementary Figure S8A). We first found that AAV-YAP^5SA^ transduction significantly increased Cyclin D1 levels in retinal explants (Fig. 8B). We next analyzed Müller cell proliferative activity through an EdU incorporation assay. In explants overexpressing YAP^5SA^, EdU labelling was strongly enhanced, with numerous patches containing high concentration of EdU-positive cells appearing within the whole tissue (Fig. 8C). Importantly, these patches coincided with regions exhibiting a high concentration of AAV-YAP^5SA^ infected cells (as assessed by Flag immunostaining) (Supplementary Figure S8B). EdU incorporation was quantified (both outside and inside these patches) and revealed a 3- to 20-fold increase in the number of positive cells compared to the control situation. Importantly, a double EdU-Sox9 labelling confirmed their Müller glial identity. Quantification revealed that up to ~25% Müller cells were proliferating in AAV-YAP^5SA^ infected explants (Fig. 8D). Together, these data reveal that YAP overactivation is sufficient to override the dormancy of murine Müller glial cells and boost their proliferative potential.

**Figure 8:**
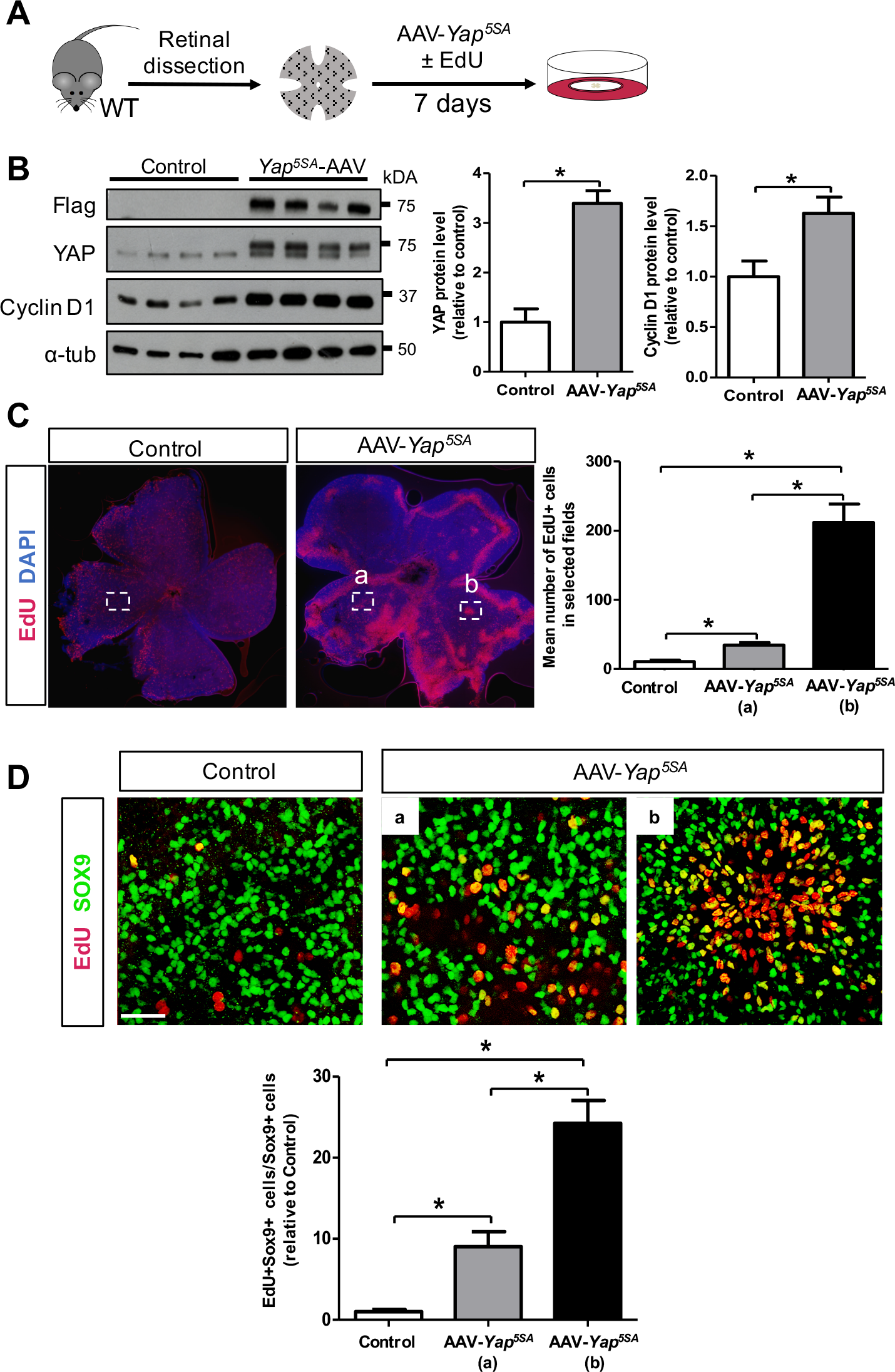
YAP overexpression in mouse Müller cells triggers their proliferative response to retinal degeneration.. **(A)** Timeline diagram of the experimental procedure used in B-D. Retinas from wild type mice were flattened, infected with AAV-*GFP* (Control) or AAV-*YAP*^*5SA*^ and cultured for 7 days. In (C, D) EdU was added to the culture medium. **(B)** Western-blot analysis of Flag-Yap^*5SA*^ and Cyclin D1 expression, *α*-tubulin labelling was used to normalize the signal. Quantification: n=4 explants for each condition. **(C)** EdU labeling on flat mounted retinas. Nuclei are counterstained with DAPI (blue). The “b” box corresponds to a region exhibiting high concentration of EdU-positive cells, while the “a” box shows an area outside such patches. Quantification: n=3 explants for each condition. **(D)** Enlargement of the explant regions delineated in C, showing EdU/Sox9 co-immunostaining (double labelled cells in yellow). Quantification: n=3 explants for each condition. In AAV-*YAP*^*5SA*^ explants, EdU^+^ (C) or EdU^+^-SOX9^+^ cells were quantified outside (a) or inside (b) the patches. Statistics: Mann-Whitney test, *p≤ 0.05. Scale bars: 500 μm (C), 20 μm (D).

## DISCUSSION

Through back and forth investigations in both mouse and *Xenopus* retinas, we discovered a pivotal role for YAP in the regulation of Müller cell response to injury (Supplementary Figure S9). We in particular reveal that YAP triggers their exit from quiescence in a degenerative context. In *Xenopus* this is accompanied with intense proliferation but not in mouse. We however demonstrate that enhancing YAP activity is sufficient to boost the naturally limited proliferative potential of mammalian Müller glia. In addition, our findings unravel a novel YAP-EGFR axis in Müller glia cell cycle re-entry that sheds a new light on the genetic network underlying their recruitment following retinal injury.

YAP knockout in several mammalian organs, such as liver, pancreas, intestine, and mammary gland, unexpectedly suggested that this factor is dispensable to maintain normal adult tissue homeostasis^40^. In line with this, we did not observe any major abnormalities in *Yap* CKO retinal morphology and function. We however revealed YAP requirement in reactive Müller glia. Reactive gliosis occurs upon retinal stress or injury^41^ and includes a series of characteristic morphological and molecular changes such as cellular hypertrophy and increased expression of intermediate filaments (nestin, vimentin and GFAP). Importantly, another feature of reactive Müller cells is also their exit from a quiescent G0 state. Although their cell cycle rarely reach S-phase, G0 to G1 progression is in particular materialized by the upregulation of genes encoding components of Cyclin D–CDK complexes, known to drive early to mid-G1 phase progression^42^. We found that many cell cycle genes, including *Ccnd* and *Cdk* genes are downregulated in *Yap* CKO reactive Müller cells, suggesting that this process is impaired in the absence of YAP. Many different transcription factors have been identified that directly regulate *Ccnd1* promoter^43^. Remarkably, YAP has been described as one of them in cancer cells^44^. In addition, *Ccnd1* was shown to be activated by YAP overexpression in the chick neural tube^45^. However, in that study, the authors reported that YAP is not required for its basal transcription. In contrast, we found in Müller cells that YAP is necessary both to maintain basal levels of Cyclin D1 in physiological conditions, and for enhancing its expression upon injury. This reinforces the hypothesis that *Ccnd1* may be a direct YAP target gene in Müller glia. This could also be the case of *Cdk6*, as previously reported in a human fibroblastic cell line^46^. Of note, although Cyclin D1 and Cyclin D2 are widely known for their role in G1-phase progression, Cyclin D3 function is still unclear. Indeed, a transcriptional regulatory activity of this Cyclin has been reported in several tissues, where it appears to promote cell cycle exit, thereby favoring differentiation^47–49^. To our knowledge, the role of Cyclin D3 in reactive Müller cells remains unknown and its link with YAP in Müller glia cell cycle reentry thus remains to be further investigated.

YAP is now well recognized as a molecular hub connecting several key signaling pathways^50^. We here reveal a cross-talk between YAP and the EGFR pathway in Müller cells following retinal degeneration. Their functional interaction was previously reported in other contexts. The EGFR ligand amphiregulin (AREG) was in particular shown to be regulated by YAP in human mammary epithelial cells or in cervical cancer cells^51,52^. We did not identify *Areg* as deregulated in *Yap* CKO retinas, but we retrieved in our RNA-Seq dataset 4 genes encoding either ligands (HBEGF and Neuregulin1), or receptors (EGFR and ERBB4) of the pathway. Interestingly, all were reported as direct YAP target genes in human ovarian cells^53^, suggesting that it could be the case in Müller cells as well. On a functional point of view, our data suggest that their decreased expression in *Yap* CKO retinas likely underlies (i) a suboptimal activity of the EGFR pathway (revealed by decreased levels of ERK1/2 and AKT phosphorylation) and (ii) the consequent failure of mutated Müller cells to upregulate G1-phase genes (as inferred from Cyclin D1 expression rescue in HB-EGF-treated *Yap* CKO retinas). Considering that EGFR signaling is a key pathway inducing Müller glia cell cycle re-entry^26–31,54,55^, such functional interaction brings YAP at the core of Müller cell reactivation mechanisms. Altogether, we propose the YAP-EGFR axis as a central player in Müller glia response to retinal damage (Supplementary Figure S9). Interestingly, this is reminiscent of the intestinal regeneration situation, where YAP-dependent EGFR signaling has previously been reported to drive tissue repair upon injury^17^.

We demonstrated that in *Xenopus*, YAP is required for Müller cell proliferation in a lesioned or degenerative context. The importance of YAP in *Xenopus* tissue repair had previously been reported in the context of epimorphic limb and tail regeneration^37,56^. Besides, we recently discovered that YAP is necessary for proper proliferation of retinal stem cells of the ciliary marginal zone, through its role in the regulation of DNA replication timing program^35^. Whether such function is conserved in proliferating Müller cells remains however to be explored. Besides, several signaling pathways, such as Notch, Wnt or Shh, were shown to regulate Müller cell proliferative response in animal models harboring retinal regeneration properties such as zebrafish or chick^2,3,57^. Given the known interplay between YAP and these pathways in various cellular contexts^11,58^, it would be interesting to seek for their potential cross-talks with YAP during Müller cell reactivation.

YAP overexpression or Hippo pathway inhibition was already demonstrated to stimulate regeneration of several injured mammalian organs, such as the heart, liver, or intestine^59–61^. Furthermore, it has recently been discovered that YAP/TAZ can act as reprogramming factors, able to turn differentiated cells into their corresponding somatic stem cells^62^. We here report that enhancing YAP activity awakes quiescent Müller cells and powerfully boosts their proliferative properties. Whether such increased proliferation leads to the production of new neuronal cells will be an important point to investigate. Noteworthy, a recent study revealed that mouse Müller cells can indeed generate new rod photoreceptors provided canonical Wnt pathway activation (to stimulate proliferation), followed by overexpression of transcription factors known to promote rod cell specification^63^. It would thus be interesting to assess in this context potential synergistic effects of YAP on the regeneration efficiency and on the consequent restoration of vision function. As a whole, by identifying YAP as a powerful inducer of Müller glia proliferation, our findings open new avenues for researches aimed at developing therapeutic strategies based on endogenous repair of the retina.

## MATERIAL AND METHODS

### Ethics statement

All animal experiments have been carried out in accordance with the European Community Council Directive of 22 September 2010 (2010/63/EEC). All animal cares and experimentations were conducted in accordance with institutional guidelines, under the institutional license D 91-272-105 for mice and the institutional license C 91-471-102 for *Xenopus*. The study protocols were approved by the institutional animal care committee CEEA n°59 and received an authorization by the “Ministère de l’Education Nationale, de l’Enseignement Supérieur et de la Recherche” under the reference APAFIS#1018-2016072611404304v1 for mice experiments and APAFIS#998-2015062510022908v2 for *Xenopus* experiments.

### Mice

Mice were kept at 21°C, under a 12-hour light/12-hour dark cycle, with food and water supplied *ad libitum*. *Yap*^*flox/flox*^ mice were obtained from Jeff Wrana’s lab^19^ and crossed with heterozygous *Rax-CreER^T2^* knock-in mice or double heterozygous *Rax-CreER^T2^*, *R26-CAG-lox-stop-lox-TdTom (Ai9)* mice from Seth Blackshaw’s lab^20^ to generate *Yap*^*flox/flox*^;*Rax-CreER^T2^* and *Yap*^*flox/flox*^;*Rax-CreER*^*T2*^;*Ai9* mice. Primer sequences used for genotyping tail snip genomic DNA are provided in Supplementary Table 1. Cre activity was induced through a single intraperitoneal injection of 4-hydroxytamoxifen (4-OHT; 1 mg) at P10, as previously described^20^. To generate the *Yap*^*flox/flox*^;*Rax-CreER^T2^;rd10* line (*Yap CKO; rd10*), *Yap*^*flox/flox*^;*Rax-CreER^T2^* mice were crossed with homozygous *rd10* mice (*Pde6b*^*rd10*^; a model of retinitis pigmentosa, with a mutation in *phosphodiesterase-6b* (*Pde6b*) gene^24^; The Jackson Laboratory, Bar Harbor, ME, USA). Chemically-induced retinal degeneration was performed through a single intra-peritoneal injection of 1-Methyl-1-nitrosourea (MNU, Trinova Biochem) at 60 mg/kg body weight, as previously described^18^.

### Xenopus

*Xenopus laevis* tadpoles were obtained by conventional procedures of *in vitro* or natural fertilization, staged according to Nieuwkoop and Faber method^64^ and raised at 18-20°C. Heat-shock-inducible dominant-negative YAP transgenic line *Tg(hsp70:dnYap-GFP, cryga:tdTomato)*^56^ (*Tg(dnYAP)*) was obtained from the UK *Xenopus* resource center (EXRC). Heat-shock-mediated dnYAP expression was induced by shifting the animals from 18-20°C to 34°C for 30 min as previously described^56^. The transgenic *Xenopus* line *Tg(Rho:GFP-NTR)* (*Tg(NTR)*) used for conditional rod ablation was generated in the lab and previously described^4^. Photoreceptor degeneration was induced by bathing the froglets in a 10 mM MTZ (Sigma Aldrich) solution for 1 week^4^. Retinal mechanical injury was performed as previously described^4^, by poking the retina once under a stereomicroscope with a needle (Austerlitz Insect Pins, 0.2 mm), without damaging the cornea or the lens. For cell proliferation assays, *Xenopus* tadpoles (*i.e.* pre-metamorphic individuals) or froglets (*i.e.* post-metamorphic individuals) were immersed in a solution containing 1 mM BrdU (5’-bromo-2’-deoxyuridine, Roche) for 3 days or 1 week, as indicated. The solution was renewed every other day.

### Mouse retinal explants

Retinas from enucleated P30 eyes were dissected in Hanks’ Balanced Salt solution (Gibco) by removing the anterior segment, vitreous body, sclera and RPE. They were then flat-mounted onto a microporous membrane (13 mm in diameter; Merk Millipore) in a twelve-well culture plate, with the ganglion cell layer facing upwards. Each well contained 700 μL of culture medium, consisting in DMEM-Glutamax (Gibco) supplemented with 1% FBS, 0.6% D-Glucose, 0.2% NaHCO_3_, 5mM HEPES, 1X B27, 1X N2, 1X Penicillin-Streptomycin. 100 ng/mL human recombinant HB-EGF (R&D systems) or vehicle was added to the culture medium from the beginning of the explant culture. Explants were maintained at 37°C in a humidified incubator with 5% CO_2_. Half of the culture medium was changed daily. For proliferation assays, 20 mM EdU was applied 1 week before fixation.

### AAV production and retinal transduction

Human YAP^5SA^ cDNA was amplified by PCR from pCMV-flag YAP2^5SA^ plasmid (a gift from Sigolène Meilhac, Addgene#27371) and subcloned into an AAV transfer plasmid, where the expression is driven by the minimal cytomegalovirus (CMV) promoter^39^. AAV vectors were produced as previously described using the co-transfection method and purified by iodixanol gradient ultracentrifugation^65^. AAV vector stocks were tittered by qPCR using SYBR Green (Thermo Fischer Scientific) as described previously^66^. The previously engineered AAV-*GFP*^39^ was used as a control. 10^11^ gc of AAV-*GFP* or AAV-*Yap*^*5SA*^ were applied on mouse retinal explants for viral transduction. Infected explants were cultured for one week before further western blot analysis or EdU incorporation assay followed by immunolabelling.

### Electroretinography

Electroretinograms (ERGs) were recorded using a Micron IV focal ERG system (Phoenix Research Labs). Mice were dark-adapted overnight and prepared for recording in darkness under dim-red illumination. They were anesthetized through intraperitoneal injection of ketamine (90 mg/ kg, Merial) and xylazine (8 mg/kg, Bayer), and were topically administered tropicamide (0.5%) and phenylephrine (2.5%) for pupillary dilation. Flash ERG recordings were obtained from one eye. ERG responses were recorded using increasing light intensities ranging from −1.7 to 2.2 log cd˙s/m^2^ under dark-adapted conditions, and from −0.5 to 2.8 log cd˙s/m^2^ under a background light that saturates rod function. The interval between flashes varied from 0.7 s at the lowest stimulus strength to 15 s at the highest one. Five to thirty responses were averaged depending on flash intensity. Analysis of a-wave and b-wave amplitudes was performed using LabScribeERG software. The a-wave amplitude was measured from the baseline to the negative peak and the b-wave was measured from the baseline to the maximum positive peak.

### Western-blotting

Each protein extract was obtained from a single retina, quickly isolated and frozen at −80°C. Retinas were then lysed in P300 buffer (20 mM Na_2_HPO_4_; 250 mM NaCl; 30 mM NaPPi; 0.1% Nonidet P-40; 5 mM EDTA; 5mM DTT) supplemented with protease inhibitor cocktail (Sigma-Aldrich). Lysate concentration was determined using the Lowry protein assay kit (DC Protein Assay; Bio-Rad) and 20 μg/lane of each sample were loaded for western-blot analysis. At least 3 individuals were tested per condition. Primary and secondary antibodies are listed in Supplementary Table 2. Protein detection was performed using an enhanced chemiluminescence kit (Bio-Rad). Each sample was probed once with anti-tubulin antibody for normalization. Quantification was done using Fiji software (National Institutes of Health)^67^.

### Whole transcriptome sequencing (RNA-Seq) and data analysis

Whole transcriptome analysis was performed on three independent biological replicates from non-injected WT, MNU-injected control and MNU-injected *Yap* CKO retinas at P60. After harvesting, both retinas for each animal were immediately frozen. RNA was extracted using Nucleospin RNA Plus kit (Macherey-Nagel). RNA quality and quantity were evaluated using a BioAnalyzer 2100 with RNA 6000 Nano Kit (Agilent Technologies). Stranded RNA-Seq libraries were constructed from 100 ng of high-quality total RNA (RIN > 8) using the TruSeq Stranded mRNA Library Preparation Kit (Illumina). Paired-end sequencing of 40 bases length was performed on a NextSeq 500 system (Illumina). Pass-filtered reads were mapped using HISAT2 2.1.0 and aligned to mouse reference genome GRCm38^68,69^. Count table of the gene features was obtained using FeatureCounts^70^. Normalization, differential expression analysis and FPKM (fragments per kilobase of exon per million fragments mapped) values were computed using EdgeR^71^. An FPKM filtering cutoff of 1 in at least one of the 6 samples was applied. A *p-value* of less than or equal to 0.05 was considered significant and a cutoff of a fold change of 1.5 was applied to identify differentially expressed genes. GO annotation was obtained using PANTHER classification system and pathways analysis was done using the Kyoto Encyclopedia of Genes and Genome (KEGG). Visualization of functional analysis data were done using GOplot R package^72^.

### RNA extraction and RT-qPCR

Total RNA was extracted from mouse neural retina or whole *Xenopus* tadpoles using RNeasy mini kit (Qiagen) or NucleoSpin RNA Plus kit (Macherey Nagel), respectively. RNA quantity was assessed using the NanoDrop 2000c UV-Vis spectrophotometer (Thermo Fisher Scientific). Total RNA was reverse transcribed in the presence of oligo-(dT)20 using Superscript II reagents (Thermo Fisher Scientific). For each RT-qPCR reaction, 1.5 ng of cDNA was used in triplicates on a CFX96 Real-Time PCR Detection System (Bio-Rad). Differential expression analysis was performed using the ΔΔCt method using the geometric mean of *Rps26, Srp72* and *Tbp* as endogenous controls^73^ for mouse genes and of *Rpl8* and *Odc1* as endogenous controls for *Xenopus* genes. For each gene, the relative expression of each sample was calculated using the mean of the controls as the reference (1 a.u.). Primers are listed in Supplementary Table 1. RT-qPCR experiments were performed on at least three mice or 3 tadpoles per condition, allowing for statistical analysis.

### Tissue sectioning, Immunofluorescence, TUNEL assay and EdU labeling

Pre- and post- metamorphic *Xenopus* individuals were anesthetized in 0.4% MS222 and then fixed in 1X PBS, 4% paraformaldehyde, overnight at 4°C. For mice, the eyes of sacrificed animals were rapidly enucleated and dissected in Hanks’ Balanced Salt solution (Gibco) to obtain posterior segment eye-cups, which were then fixed in 1X PBS, 4% paraformaldehyde for 1 h at 4°C. Fixed samples were dehydrated, embedded in paraffin and sectioned with a Microm HM 340E microtome (Thermo Scientific). Immunostaining on retinal sections or explants was performed using standard procedures with the following specificities: (i) An antigen unmasking treatment was done in boiling heat-mediated antigen retrieval buffer (10 mM sodium citrate, pH 6.0) for 20 min; (ii) For *Xenopus* sections, this was followed by a 45 min treatment in 2N HCl at room temperature; (iii) Incubation timing was increased at all steps for immunolabelling on retinal explants. All primary and secondary antibodies are listed in Supplementary Table 2. Nuclei were counterstained with 1μg/ml DAPI (Thermo Fisher Scientific) or Hoechst (Sigma). H&E staining was performed using standard procedure. Detection of apoptotic cells was conducted on retinal sections using DeadEnd Fluorometric TUNEL System (Promega) according to the manufacturer’s instructions. EdU incorporation was detected on flat mounted mouse retinal explants using the Click-iT EdU Imaging Kit (Thermo Fisher Scientific) following the manufacturer’s recommendations. Coverslips were mounted using Fluorsave Reagent (Millipore, USA).

### Imaging

Fluorescence and brightfield images were acquired using an ApoTome-equipped AxioImager.M2 microscope or a *Zeiss LSM*710 confocal microscope. Whole retinal explants were imaged using an AxioZoom.v16 (Zeiss). Images were processed using Zen (*Zeiss*), Fiji (National Institutes of Health) and Photoshop CS4 (Adobe) softwares. The same magnification, laser intensity, gain and offset settings were used across animals for any given marker.

### Quantification and statistical analysis

All quantifications were performed by manual counting except for *Xenopus* Müller cells. In that case, anti-Glutamine synthetase fluorescence intensity was measured across the entire section using Photoshop CS4 (Adobe) software and normalized using the DAPI signal. 3 retinas were used per condition. BrdU-positive cells were counted within the *Xenopus* neural retina (after exclusion of the ciliary marginal zone). 5 to 10 sections were analyzed for each retina, and an average number was calculated from at least 3 individuals. Mean number of EdU-labelled cells in mouse retinal explants were calculated from 3 different fields of 10^4^ μm^2^ per retina and using 3 explants per condition. SOX9-positive mouse Müller cells were quantified by considering one entire retinal section from three different mice for each condition. Statistical analysis was performed using the non-parametric Mann-Whitney test in all experiments except ERG, for which we applied a two-way ANOVA test. p-value ≤*0.05* was considered significant. All results are reported as mean ± SEM.

## ACKNOWLEDGMENTS

We are thankful to S. Blackshaw, J. Wrana and S. Meilhac for mouse lines and plasmids. We are grateful to E-K. Grellier and S. Lourdel for their help with the maintenance of mouse colonies. We would also like to thank E-K. Grellier for performing the ERG recordings, A. Chesneau, R. Langhe and S. Lourdel for their technical assistance with *Xenopus* fertilization experiments and immunostaining experiments. This work has benefited from the facilities and expertise of the high throughput sequencing core facility of I2BC (Centre de Recherche de Gif - http://www.i2bc-saclay.fr/). This research was supported by grants to M.P. from the FRM, Association Retina France, Fondation Valentin Haüy and the DIM Région île de France. A.H. is a Retina France association and ARC fellow, D.G.G is a FRM fellow.

## AUTHOR’S CONTRIBUTION

A.H., D.A., D.G.G., J.B. designed and performed the experiments and analyzed the data, D.D. supervised AAV production, M.L. revised the manuscript, J.R. designed the study, analyzed the data, M.P. designed the study, analyzed the data, wrote the manuscript with the help of A.H. and D.A. and supervised the study.

---

**Supplementary Figure S1:**
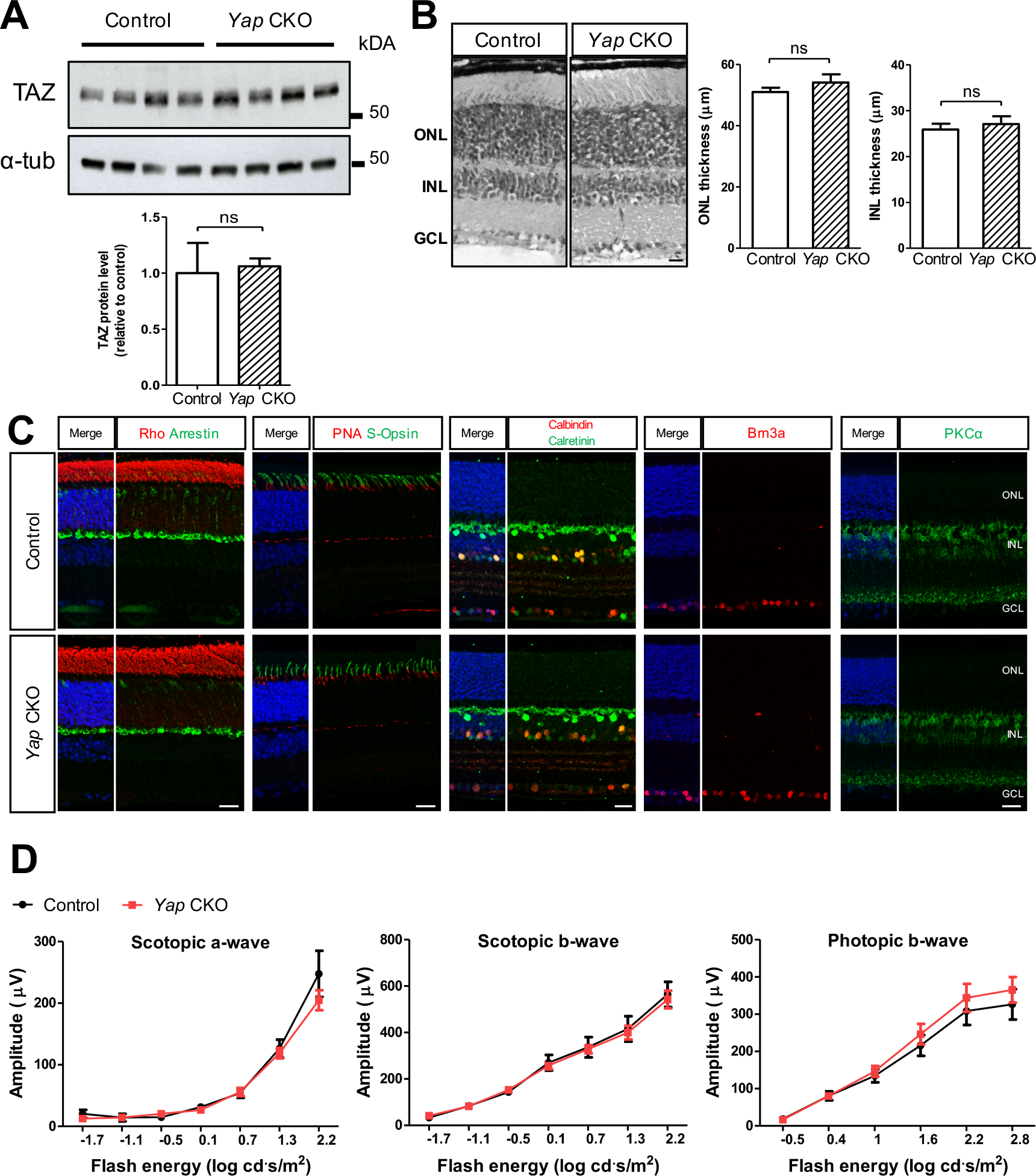
*Yap* CKO mice do not show any major retinal defects.. **(A)** Western-blot analysis of TAZ expression on retinal extracts from Control and *Yap* CKO mice injected with 4-OHT as shown in Fig. 1A. *α*-tubulin labelling was used to normalize the signal. Quantification: n=4 mice for each condition. **(B)** H&E staining on retinal sections from Control and *Yap* CKO mice injected with 4-OHT as shown in Fig. 1A. Quantification of INL and ONL thickness in the central retina: n=4 mice for each condition. **(C)** Retinal sections from Control and *Yap* CKO mice injected with 4-OHT as shown in Fig. 1A, immunostained as indicated for a rod marker (Rhodopsin, Rho), cone photoreceptor markers (Arrestin, lectin PNA and S-Opsin), a horizontal cell marker (Calbindin), an amacrine cell marker (Calretinin), a ganglion cell marker (Brn3a) and a bipolar cell marker (PKCα). Nuclei are counterstained with DAPI (blue). **(D)** ERG-derived intensity response curves of the average scotopic a-wave, scotopic b-wave and photopic b-wave, in Control (n=6) and *Yap* CKO (n=9) mice injected with 4-OHT as shown in Fig. 1A. Two-way ANOVA tests revealed no significant differences. ONL: outer nuclear layer; INL: inner nuclear layer; GCL: ganglion cell layer. Statistics: Mann-Whitney test (A, B), ns: non-significant. Scale bars: 50 μm (B), 20 μm (C).

**Supplementary Figure S2:**
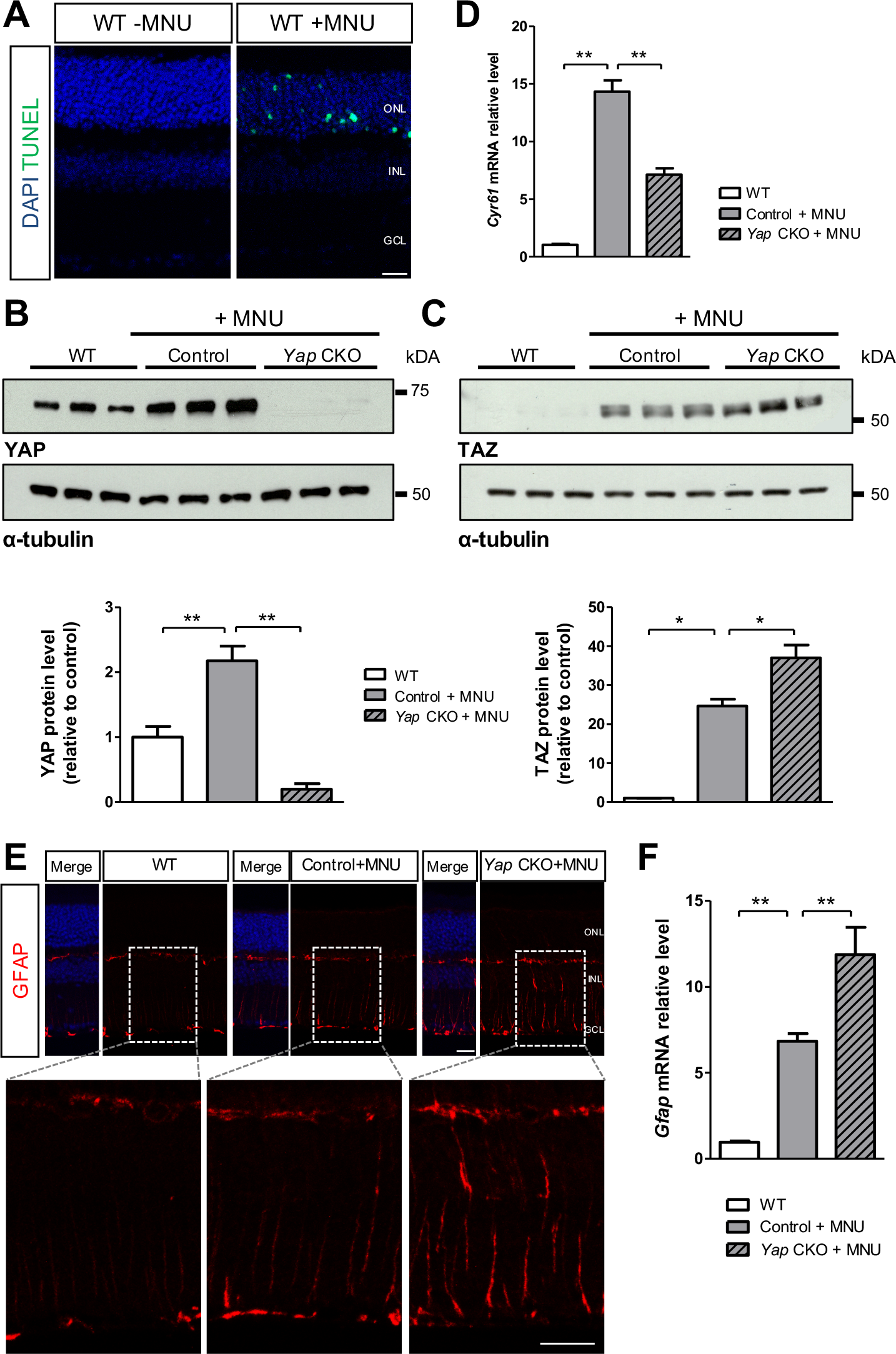
YAP/TAZ expression in *Yap* CKO retinas following MNU-induced degeneration.. **(A)** TUNEL assay on retinal sections from non-injected and MNU-injected WT mice (16 hours post-injection). Nuclei are counterstained with DAPI (blue). **(B, C)** Western-blot analysis of YAP and TAZ expression on retinal extracts from non-injected WT mice or from Control and *Yap* CKO mice injected with 4-OHT and MNU as shown in Fig. 1F. *α*-tubulin labelling was used to normalize the signal. Quantification: n=3 mice for each condition. **(D)** RT-qPCR analysis of *Cyr61* expression on the same experimental conditions (at least 5 biological replicates per condition were used). **(E)** Retinal sections immunostained for GFAP. Nuclei are counterstained with DAPI (blue). The regions delineated with dashed lines are enlarged in the lower panels. **(F)** RT-qPCR analysis of *Gfap* expression (at least 5 biological replicates per condition were used). ONL: outer nuclear layer; INL: inner nuclear layer; GCL: ganglion cell layer. Statistics: Mann-Whitney test, *p≤ 0.05, ** p≤ 0.01. Scale bar: 20 μm.

**Supplementary Figure S3.**
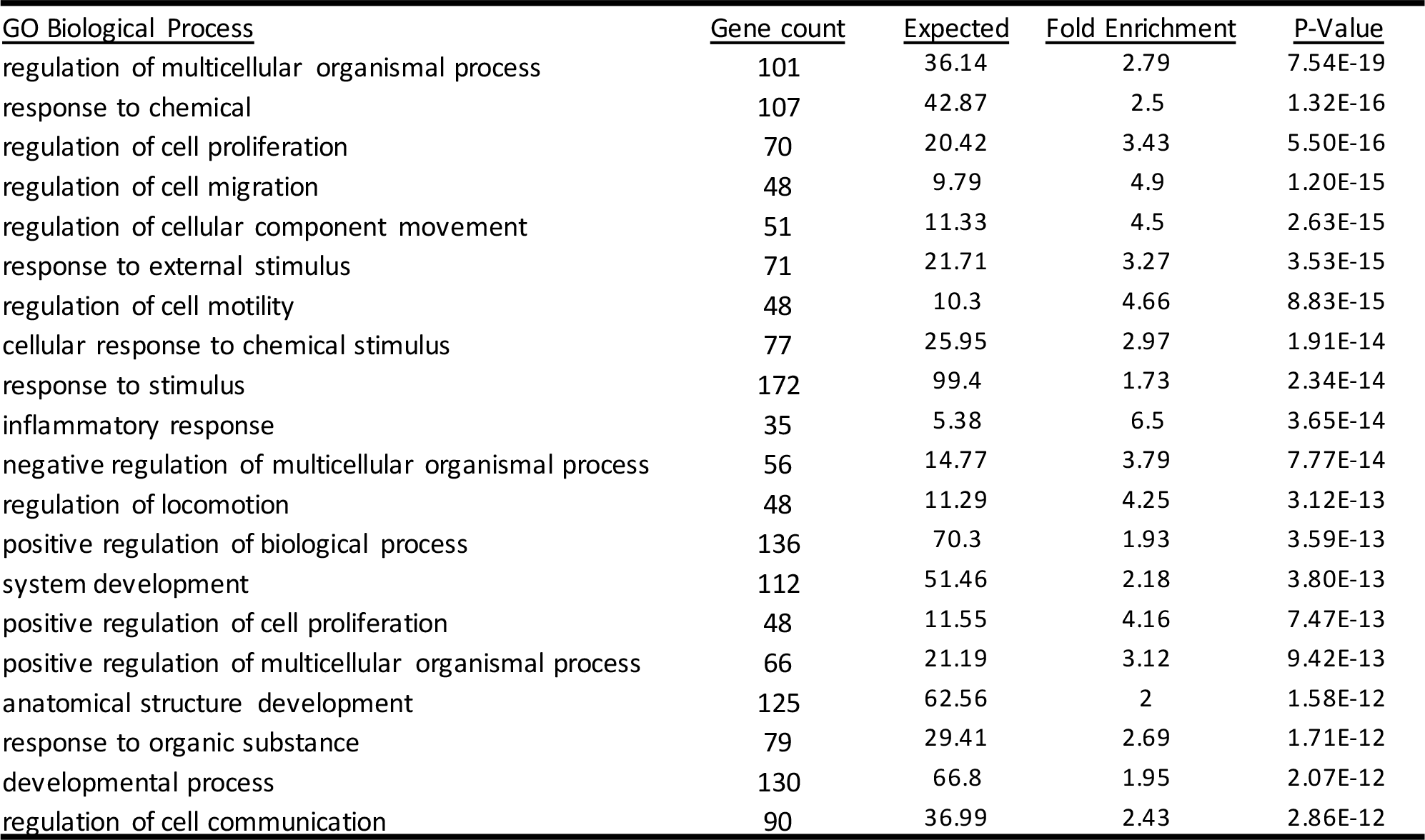
Deregulated biological processes in *Yap* CKO retinas responding to MNU-induced degeneration.. List of the top twenty over-represented GO Biological Processes to which belong the 305 DEGs retrieved from our RNAseq dataset comparing MNU-injected Control and MNU-injected *Yap* CKO mice.

**Supplementary Figure S4:**
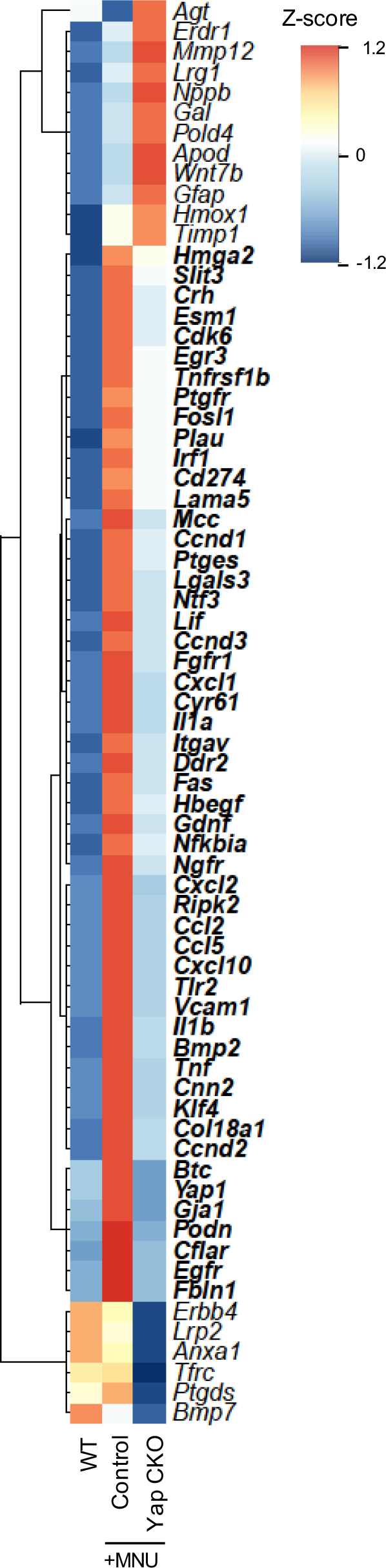
Deregulated genes related to cell proliferation in *Yap* CKO mice responding to MNU-induced degeneration.. Hierarchical clustering analysis of the 70 DEGs that belong to the GO Biological Process “regulation of cell proliferation” (GO:0042127). The resulting heatmap represents z-score expression values for each condition. Genes in bold are those that are (*i*) expressed at very low levels in non-injected WT mice, (*ii*) strongly upregulated in MNU-injected Control mice, (*iii*) only moderately enriched in MNU-injected *Yap* CKO mice.

**Supplementary Figure S5:**
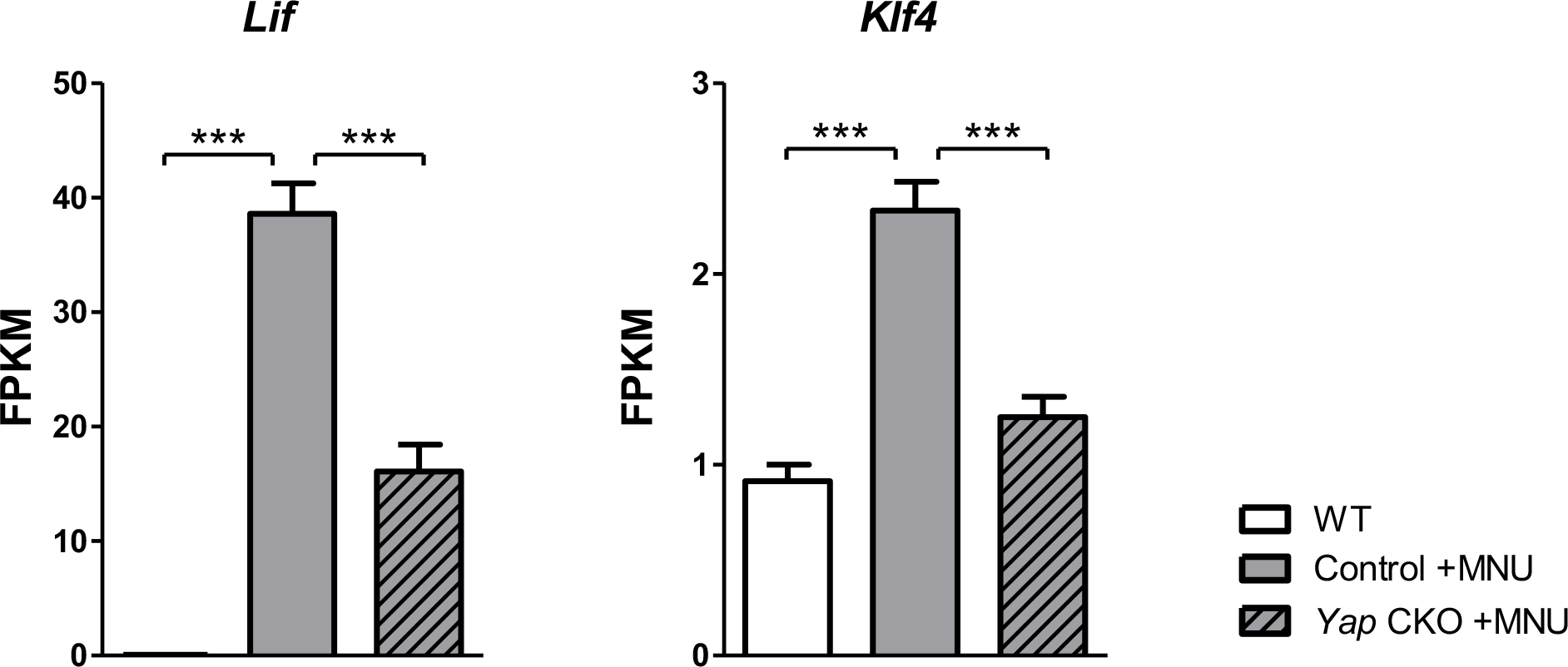
Pluripotent/reprograming gene upregulation in response to MNU injection is compromised in *Yap* CKO mice.. Relative RNA expression (in FPKM; data retrieved from the RNA-seq experiment) of *Klf4 and Lif*, in retinas from non-injected WT mice or Control and *Yap* CKO mice injected with 4-OHT and MNU as shown in Figure 1F. Statistics: P-values were obtained using EdgeR, *** p≤ 0.001.

**Supplementary Figure S6:**
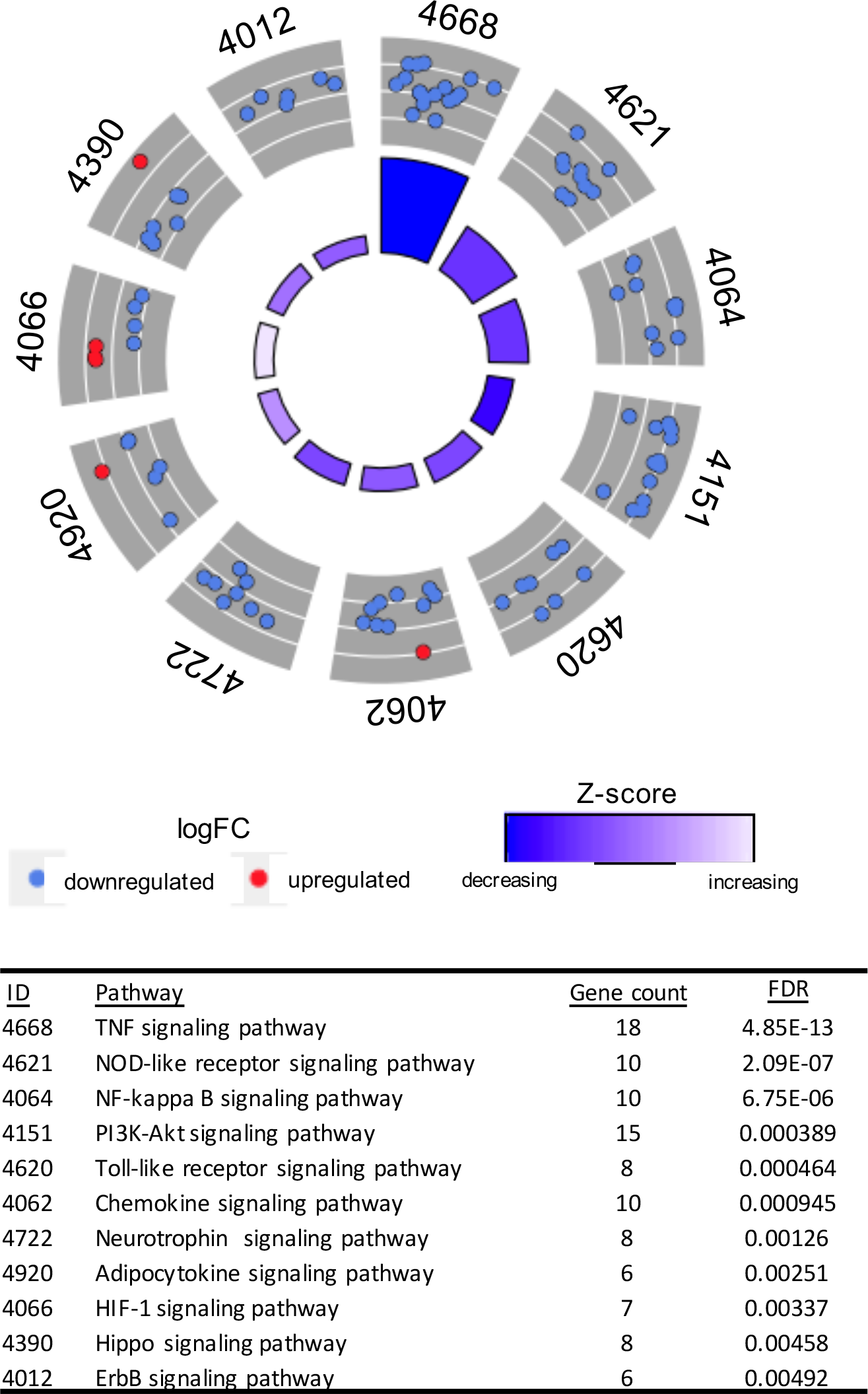
Deregulated signaling pathways in *Yap* CKO retinas responding to MNU-induced degeneration.. Pathway analysis of the RNA-seq dataset comparing non-injected WT mice or MNU-injected Control and *Yap* CKO mice. The eleven most deregulated signaling pathways identified by Kyoto Encyclopedia of Genes and Genome (KEGG) enrichment analysis are shown. Downregulated (blue dots) and upregulated (red dots) genes within each signaling pathway are plotted based on logFC. Z-score bars indicate if an entire signaling pathway is more likely to be increased or decreased based on the behavior of genes it contains. The table shows pathway identity number (ID) depicted on the circle, associated pathway name, gene count and False Discovery Rate (FDR).

**Supplementary Figure S7:**
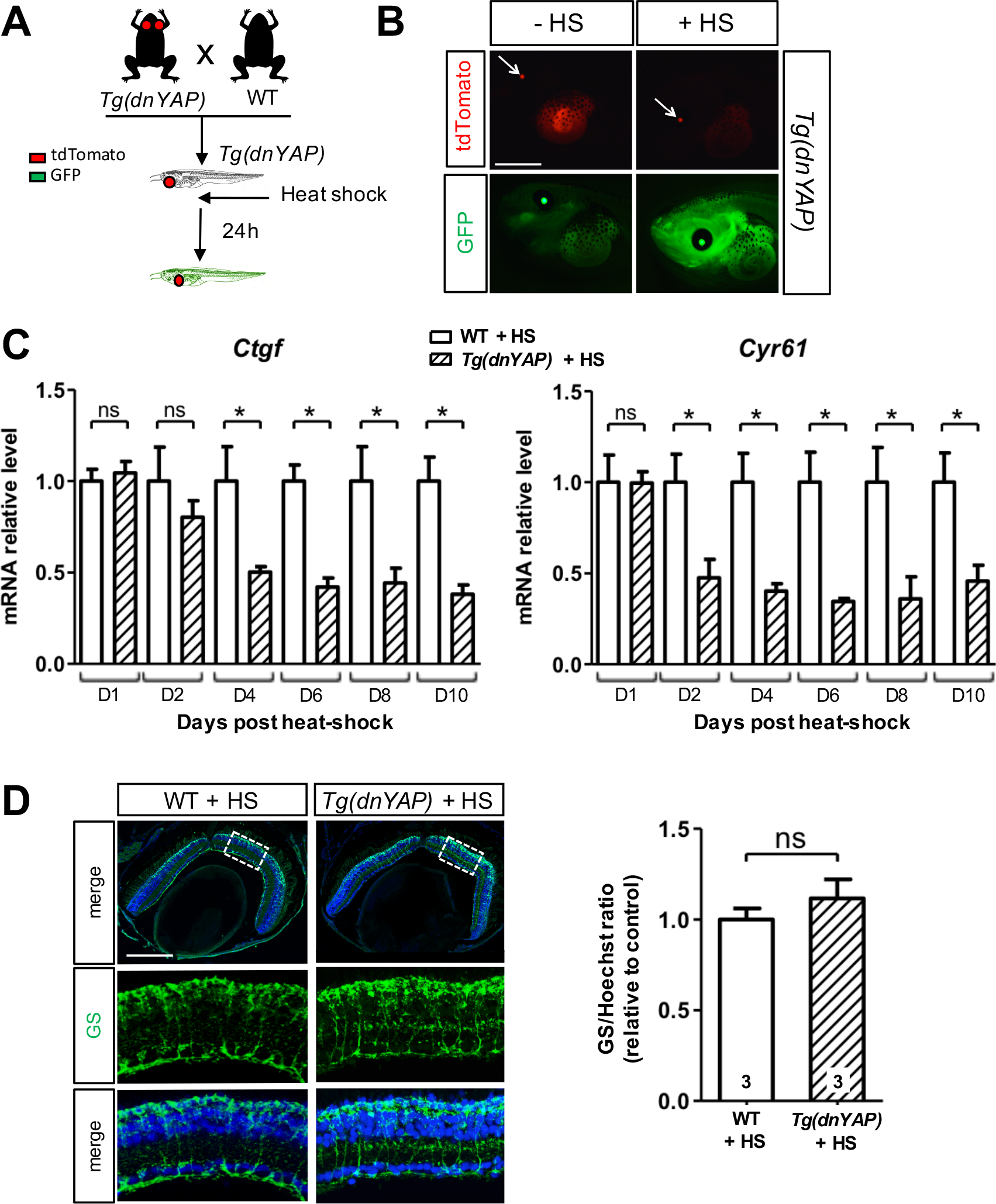
Validation of the *Xenopus Tg(dnYAP)* line.. (**A**) Timeline diagram of the experimental procedures used in B-D. Wild type (WT) *Xenopus* were crossed with *Tg(dnYAP)* individuals. *Tg(dnYAP)* tadpoles were selected within the progeny, heat-shocked and harvested 24 hours later. (**B**) Images of tadpole anterior region. Presence of the transgene is assessed by tdTomato lens expression (red, arrows). Of note, tdTomato is very bright and its fluorescence can also be seen in the GFP channel. Heat shock effectiveness is assessed by GFP expression in the entire tadpole. (**C**) RT-qPCR analysis of two YAP target genes, *Ctgf* and *Cyr61*, in tadpoles harvested 1 to 10 days after the heat shock (3 biological replicates per condition). For each time point, the control transcript level is set to 1. (**D**) Retinal sections from heat-shocked WT or *Tg(dnYAP)* tadpoles, immunostained for glutamine synthetase (GS, green). Nuclei are counterstained with Hoechst (blue). The regions delineated with dotted lines are enlarged in the lower panels. For quantification, the intensity of the green fluorescence signal (GS) was measured and normalized against the blue fluorescence signal (Hoechst) of the same retina. The number of animals tested for each condition is indicated on the corresponding bar. Statistics: Mann-Whitney test, *p≤ 0.05; ns: non-significant. Scale bars: 1 mm (B), 50 μm (D), 200 μm (E).

**Supplementary Figure S8:**
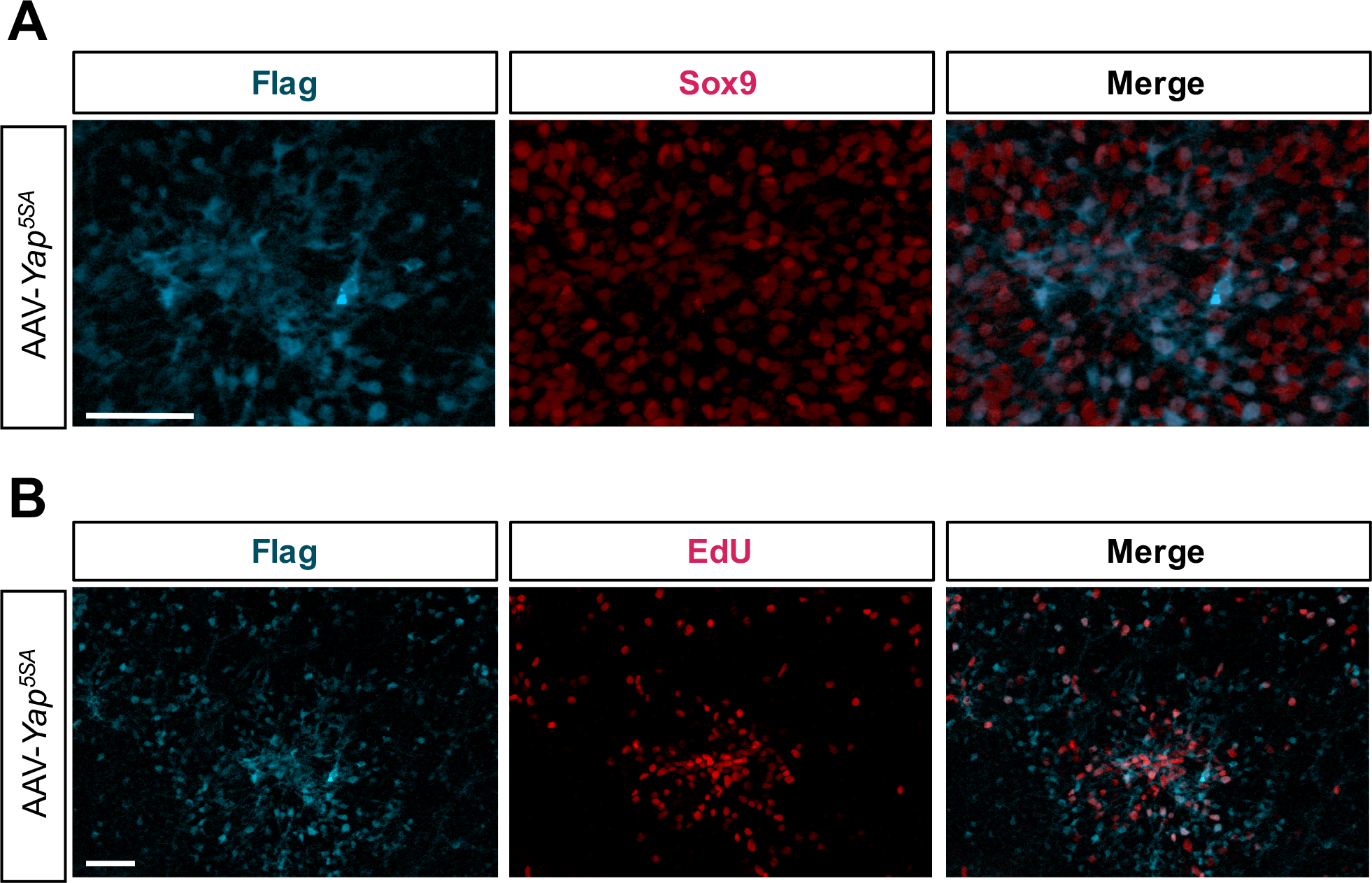
YAP overexpression in mouse Müller cells triggers their proliferative response to retinal degeneration. **(A, B)** Flag-Yap^*5SyA*^ and Sox9 or EdU labeling on flat mounted retinal explants infected with AAV-*YAP*^*5SA*^ as in Fig. 8A. Scale bars: 50 μm.

**Supplementary Figure S9:**
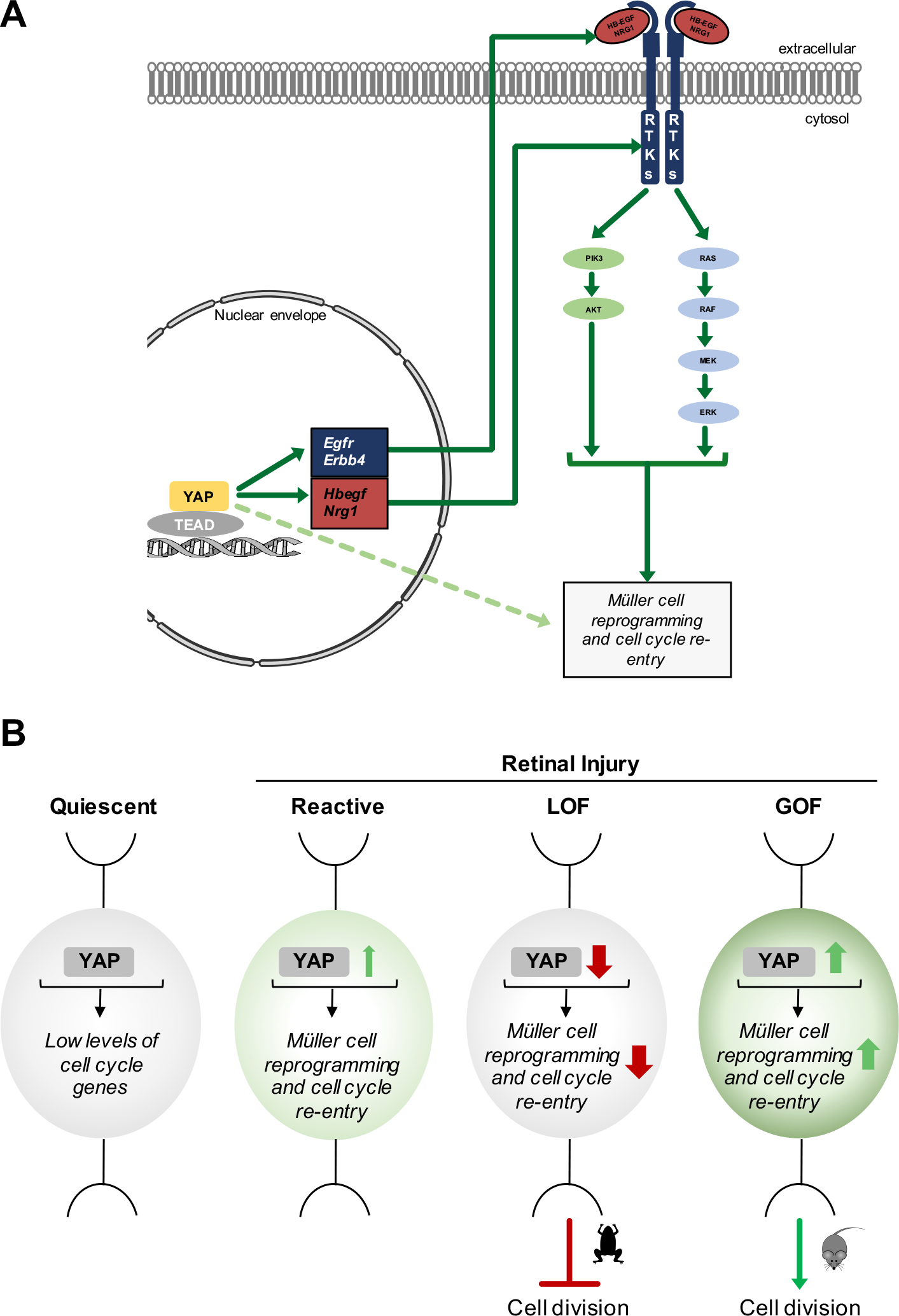
Proposed model of YAP-EGFR pathway interaction in Müller glia cell cycle re-entry upon injury.. **(A)** Our results support the idea that YAP and EGFR signaling functionally interact in a retinal degenerative context to induce cell cycle genes in Müller glia. YAP likely acts upstream the EGFR pathway by regulating the expression of EGFR receptors (EGFR and ERBB4) and ligands (HBEGF and NRG1). The EGFR pathway would in turn activate Müller cell reprograming and cell cycle re-entry, as previously reported^26–30^. Of note, YAP may also independently impact these processes through direct regulation of cell cycle and reprogramming genes, such as those identified in our RNAseq dataset (dashed arrow). **(B)** Schematics summarizing our data. In a quiescent Müller cell (grey), YAP expression maintains a basal level of cell cycle genes. Upon retinal degeneration, YAP level rises in reactive Müller cells (light green), which triggers the upregulation of reprograming and cell cycle genes. YAP loss of function (LOF) impairs Müller cell reprograming and cell cycle re-entry. In *Xenopus* which is endowed with regenerative properties, this prevents Müller cell proliferation. In mouse, YAP gain of function (GOF) is sufficient to enhance cell cycle gene expression levels (dark green) and to trigger Müller cell proliferation.

**Supplementary Table 1:**
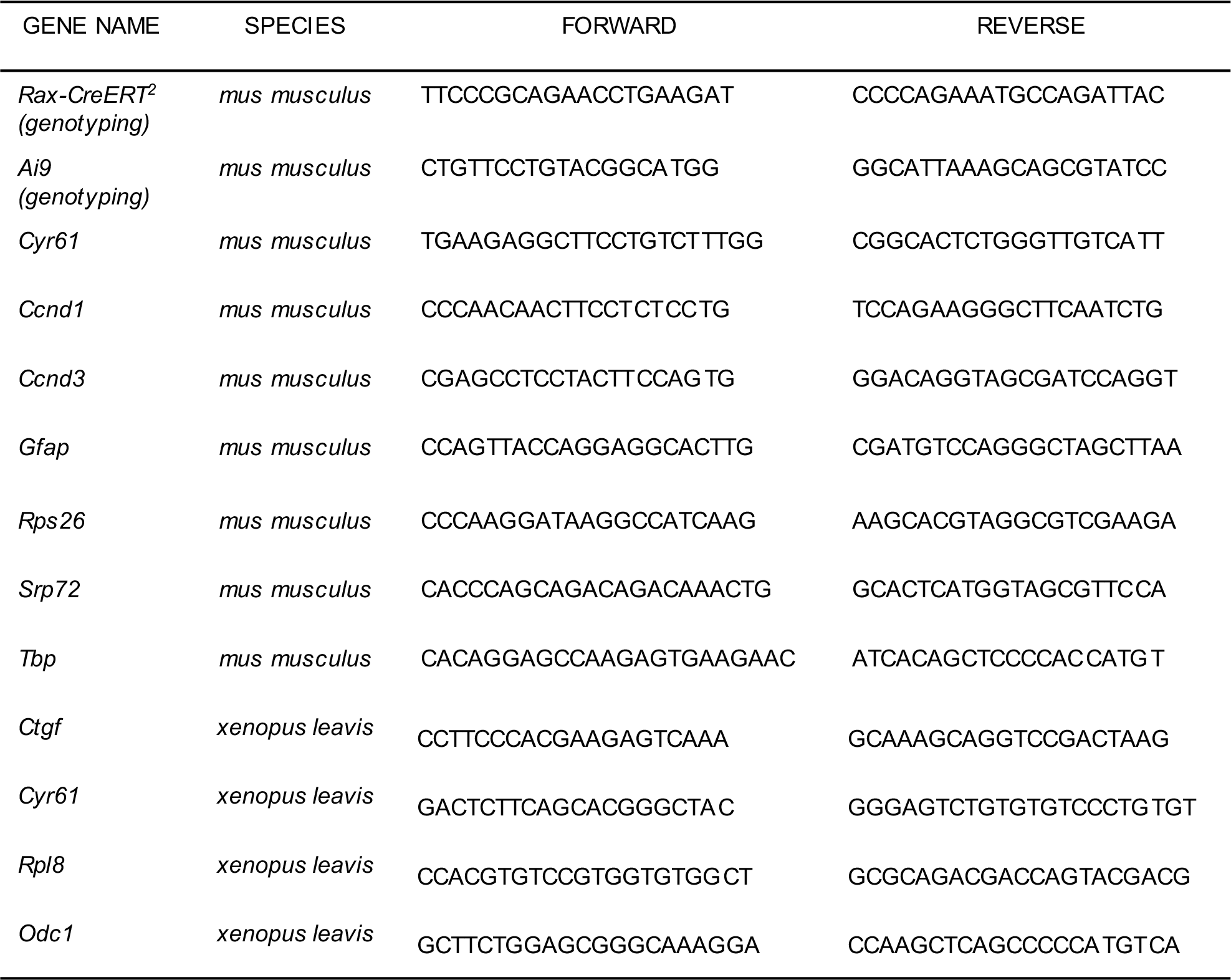
List of primers.

**Supplementary Table 2:** List of antibodies.

|  |  |  |  | Experiments |  | Experiments |
| --- | --- | --- | --- | --- | --- | --- |
|  |  |  |  | on mice |  | on xenopus |
| Antigene | Host | Supplier | Reference | Dilution (IHC) | Dilution (WB) | Dilution (IHC) |
| Primary antibodies |  |  |  |  |  |  |
| AKT | mouse | SIGMA | P2482 |  | 1:1,000 |  |
| Alpha-tubulin | mouse | SIGMA | T5168 |  | 1:10,000 |  |
| Alpha-tubulin | mouse | SIGMA | T9026 |  | 1: 5,000 |  |
| Arrestin | rabbit | EMD Millipore | AB15282 | 1:1,000 |  |  |
| Brn3a | mouse | Santa Cruz | sc-8429 | 1:100 |  |  |
| BrdU |  |  |  |  |  | 1:200 |
| Calbindin D-28k | rabbit | Swant | 300 | 1:100 |  |  |
| Calretinin | mouse | EMD Millipore | MAB1568 | 1:1,000 |  |  |
| Cyclin D1 | rabbit | Abcam | ab134175 | 1:100 | 1:1,000 |  |
| Cyclin D3 | mouse | Cell Signaling | 2936 | 1:100 | 1:1,000 |  |
| ERK1/2 (p44/42 MAPK) | rabbit | Cell Signaling | 9102 |  | 1:1,000 |  |
| Flag | mouse | Sigma | F3165 | 1:300 |  |  |
| Flag | rabbit | Sigma | F7425 |  | 1:1000 |  |
| GFAP | rabbit | Dako | Z0334 | 1:500 |  |  |
| Glutamine synthetase | mouse | Abcam | Ab64613 |  |  | 1:200 |
| Lectin PNA alexa 568 conjugate | peanut | Thermo Fisher Scientific | L32458 | 1:500 |  |  |
| P-AKT (Ser473) | rabbit | Cell Signaling | 9271 |  | 1:1,000 |  |
| P-ERK1/2 (Thr202/Tyr204) | rabbit | Cell Signaling | 4370 | 1:100 | 1:1,000 |  |
| Red Fluorescence Protein | rabbit | Rockland | 600-401-379 | 1:100 |  |  |
| Rhodopsin | mouse | EMD Millipore | MAB5316 | 1:2,000 |  |  |
| Rhodopsin | mouse | EMD Millipore | MABN15 |  |  | 1:500 |
| S-opsin | rabbit | EMD Millipore | AB5407 | 1:500 |  |  |
| Sox9 | rabbit | EMD Millipore | AB5535 | 1:300 |  |  |
| TAZ | rabbit | Abcam | ab110239 |  | 1:1,000 |  |
| Vimentin | rabbit | Santa Cruz | sc-7557 | 1:200 |  |  |
| YAP | mouse | Abcam | ab56701 | 1:50 | 1:1,000 |  |
| YAP | rabbit | Abcam | ab52771 |  |  | 1:100 |
| Secondary antibodies |  |  |  |  |  |  |
| Alexa 488 anti-mouse IgG1 | goat | Thermo Fisher Scientific | A21121 | 1:500 |  |  |
| Alexa 488 anti-rabbit | donkey | Thermo Fisher Scientific | A21206 | 1:500 |  |  |
| Alexa 555 anti-mouse IgG1 | goat | Thermo Fisher Scientific | A21127 | 1:500 |  |  |
| Alexa 555 anti-mouse IgG2b | goat | Thermo Fisher Scientific | A21147 | 1:500 |  |  |
| Alexa 555 anti-mouse IgG2a | goat | Thermo Fisher Scientific | A21127 | 1:500 |  |  |
| Alexa 555 anti-rabbit | donkey | Thermo Fisher Scientific | A31572 | 1:500 |  |  |
| Alexa 488 anti-mouse | goat | Thermo Fisher Scientific | A11001 |  |  | 1:1,000 |
| Alexa 594 anti-mouse | goat | Thermo Fisher Scientific | A11005 |  |  | 1:1,000 |
| Alexa 594 anti-rat | goat | Thermo Fisher Scientific | A11007 |  |  | 1:1,000 |
| Alexa 488 anti-rabbit | goat | Thermo Fisher Scientific | A11008 |  |  | 1:1,000 |
| Alexa 594 anti-rabbit | goat | Thermo Fisher Scientific | A11012 |  |  | 1:1,000 |
| HRP anti-mouse IgG | goat | Sigma-Aldrich | A4416 |  | 1:5,000 |  |
| HRP anti-rabbit IgG | donkey | GE Health | NA934V |  | 1:5,000 |  |

